# Keratin 6a attenuates Toll-like receptor-triggered proinflammatory response in corneal epithelial cells by suppressing ELKS/IKKε-dependent activation of NF-κB

**DOI:** 10.1101/2023.10.24.563888

**Authors:** Jonathan K. Chan, Yan Sun, Anand Bhushan, Belinda Willard, Connie Tam

## Abstract

The corneal epithelium at the ocular surface is constantly exposed to the environment and represents the first line of defense against infection, mechanical injury or chemical irritation. Through TLR-mediated recognition of pathogen- and damage-associated molecular patterns, it engages in direct antimicrobial responses and alerts the immune system on intruder and tissue damage by secreting pro-inflammatory and chemotactic cytokines that promote immune cell infiltration. How the corneal epithelium downregulates TLR signaling is unclear, yet it highly expresses keratin 6a (K6a), a cytoskeletal protein that has emerged to play essential regulatory roles in corneal innate immune response. Here we report that mice harboring genetic deletion of K6a are more susceptible to developing bacterial keratitis with unresolved corneal opacification and higher bacterial load. Such disease phenotype is caused by the increased pro-inflammatory cytokine and chemokine secretions from the K6a-null corneal epithelium, which further promotes the infiltration of immune cells and their associated pro-inflammatory response. Using human corneal epithelial cells immortalized by telomerase reverse transcriptase (hTCEpi cells), we demonstrated that knocking down K6a enhances NF-κB/ RelA-dependent cytokine and chemokine expression. Moreover, proteomic screen reveals that K6a interacts with ELKS, a critical NEMO-binding scaffold that links between canonical IKKα/β and the principal cytoplasmic inhibitor of RelA, i.e. IκBα., to promote its phosphorylation and degradation. Surprisingly, K6a does not antagonize any of these canonical NF-κB signaling events. Instead, we found that ELKS in addition to canonical IKKs interacts with the atypical IKK member IKKε. Furthermore, knockdown of K6a in hTCEpi cells promotes ELKS-dependent phosphoactivation of IKKε, which in turn phosphorylates and activates RelA. Our study thus demonstrated an unexpected role of cytosolic K6a as a novel negative regulator of TLR/NF-κB signaling in preventing excess proinflammatory cytokine and chemokine expressions. It further highlighted the functional importance of ELKS as a common signaling scaffold for both canonical and atypical IKK-dependent activation of NF-κB in corneal epithelial cells. Using both IKK classes other than only canonical IKKs for TLR/NF-κB induction as in other cell types including myeloid immune cells suggest that the cornea epithelium is more flexible in modulating its inflammatory response, which could greatly minimize corneal damage while preserving its essential functions for barrier protection and light refraction.

## Introduction

The cornea is a dome-shaped avascular tissue at the front of the eye which serves as a protective barrier and provides refractive power for light focusing onto the retina (de Paiva, St Leger, and Caspi 2022). Located at the outmost region, the corneal epithelium is comprised of stratified squamous epithelial cell layers supported by a single layer of basal cells. Like in any mucosal epithelia including the gastrointestinal, respiratory and genitourinary tracts, the corneal epithelium is constantly exposed to the environment and represents the first line of defense against infection, mechanical injury or chemical irritation (de Paiva, St Leger, and Caspi 2022). Its ability in sensing these threats relies in part by its membrane expression of Type I glycoproteins from the Toll-like receptor (TLR) family. These germline-encoded receptors recognize conserved signature motifs in microbes known as pathogen associated molecular patterns (Duan et al. 2022) as well as molecules released from damaged or dying cells known as damage associated molecular patterns (Piccinini and Midwood 2010). To date, thirteen TLRs have been identified based on their homologous intracellular Toll/IL-1 receptor (TIR) domain and are classified based on their cellular localization at the cell surface membrane or endosomal/ lysosomal compartment (Duan et al. 2022).

After cognate ligand binding, all TLRs except TLR3 recruit MYD88 to form a large multiprotein complex at the membrane facing the cytosol known as the Myddosomes (Pereira and Gazzinelli 2023). Assembled Myddosomes dissociate into the cytosol, where they recruit the E3 ligase TRAF6 to ubiquitinate itself and the essential NF-κB adaptor called NEMO with K63-linked polyubiquitin chains. These chains recruit and activate the MAP3K TAK1 (Walsh, Lee, and Choi 2015), which then phosphoactivates canonical IKKα/β members associated with NEMO, resulting in the eventual induction of NF-κB-dependent transcription of pro-inflammatory cytokines and chemokines. Another MAP3K lying downstream of TRAF6 that activates NF-κB is MEKK3 (Qin et al. 2006; Yao et al. 2007; Fraczek et al. 2008). Likewise, TLR3 (and also some TLR4) through interacting with the cytosolic TRIF adaptor associates with TRAF3, which auto-ubiquitinates itself with K63-linked polyubiquitin chains recruiting atypical IKK members TBK1 and IKKε (Hacker, Tseng, and Karin 2011; Lin et al. 2023). TBK1 and IKKε upon autophosphorylation phosphoactivate the transcription factor IRF3, resulting in the induction of Type I interferon response (Hemmi et al. 2004; Perry et al. 2004; Shu et al. 2013).

In freshly dissected human donor corneas or primary corneal epithelial cells, the expressions of TLR2, 3, 4, 5, 7 and 9 have been confirmed by FACS or immunohistochemistry (Song et al. 2001; Ueta et al. 2005; Li et al. 2006; Hozono et al. 2006; Wu, Gao, and Ren 2007; Jin et al. 2007; Redfern, Reins, and McDermott 2011). TLR2, 4 and 5 located on the cell surface are responsible for detecting bacterial lipoteichoic acids (LTA), lipopolysaccharides (LPS) and flagellin, while TLR3, 7, and 9 are present within the endosomes recognizing microbial dsRNA, ssRNA and unmethylated CpG motifs. The wide range of host-derived molecules serving as DAMPS for TLR sensing and released at damaged corneal epithelium is currently an active area of research (Lema, Reins, and Redfern 2018; Fowler et al. 2023) In all cases, the chemotactic cytokines and immune mediators synthesized and secreted into the extracellular milieu by the corneal epithelium are instrumental for attracting myeloid immune cells and priming their immune activities, resulting in pathogen eradication and dead cell removal (Hazlett 2004; Pearlman et al. 2008; Pearlman et al. 2013). However, aberrant corneal epithelial TLR activities are often accompanied by protracted immune cell infiltration during microbial keratitis and in autoimmune Sjöregen’s syndrome (Redfern and McDermott 2010). If not treated properly the sequelae of these inflammatory conditions can inflict collateral damage to the remaining healthy cornea tissues compromising their barrier and refractive functions, leading to extensive scarring and even vision loss. Interestingly, while TLR2/4 agonist MPLA and TLR7 agonist imiquimod are approved by the Food and Drug Administration as a vaccine adjuvant and for treating viral infection and skin cancer respectively (Wang et al. 2020; Rolfo et al. 2023), no TLR antagonists have been approved for clinical use perhaps due to systemic toxicities. As such, the development of TLR inhibitors that are cell or tissue type specific and with improved tolerability can have beneficial therapeutic potentials for unmet medical needs.

Keratins are intermediate filament proteins that are abundantly expressed in the skin and mucosal epithelium. With more than 50 members in human, keratin proteins are classified as either acidic type I (K9–K10, K12–K28, and K31–K40) or basic type II (K1–K8 and K71–K86 (Jacob et al. 2018; Redmond and Coulombe 2021). Specific members from type 1 and type 2 keratins form obligatory heterodimers that are incorporated into structural filamentous networks providing cellular tensile strength. Emerging evidence reveal that a minor portion of keratin remains as soluble within the cytosol and possesses signaling properties which regulate diverse cellular functions, including autophagy, apoptosis, cell growth and survival, innate immunity, to name a few (Jacob et al. 2018; Redmond and Coulombe 2021). In the retina, K8 is upregulated in patients with neovascular AMD which appears to provide cytoprotective function against oxidative stress-induced necrotic cell death in retinal pigment epithelial cells by inducing autophagy (Baek et al. 2017; Baek et al. 2021; Shin et al. 2022). Previously, we have elucidated that cytosolic K6a is processed within corneal epithelial cells and released antimicrobial peptides called KAMPs that have potent bactericidal activities (Tam et al. 2012; Lee et al. 2016; Chan et al. 2018). We recently have reported that KAMPS also have anti-inflammatory activities by downregulating TLR signaling (Sun et al. 2023). Specifically, we showed that KAMPS compete with bacterial ligands for binding with TLR2 and TLR2/4 co-receptors (MD2, CD14) on myeloid immune cells, promoting TLR2/4 endocytosis and reducing their cell surface availability for bacterial ligand binding. Topical KAMP treatment further effectively alleviated experimental bacterial keratitis by substantially reducing corneal opacification, inflammatory cell infiltration, and bacterial burden (Sun et al. 2023). Whether cytosolic K6a within the corneal epithelium itself possesses additional function in innate immunity regulation is unknown.

Using mice harboring genetic deletion of keratin 6a in keratitis models, we here discovered that within the corneal epithelium cytosolic K6a suppresses TLR-induced expression of proinflammatory cytokines and chemokines independent of its innate immune properties as KAMPS. Mechanistic studies using human corneal epithelial cells immortalized by telomerase reverse transcriptase (hTCEpi cells) revealed that the anti-inflammatory function of cytosolic K6a is mediated through its interaction with the NEMO-associated scaffold ELKS but does not suppress ELKS-dependent canonical IKKα/β phosphorylation of IκBα. Instead, ELKS interacts with atypical IKKε while K6a antagonizes both the activation of IKKε and its phosphorylation of NF-κB/RelA. We further demonstrated that K6a’s anti-inflammatory action within the corneal epithelium serves to restrain the influx of myeloid immune cells that contribute to corneal inflammatory pathology in both sterile and bacterial keratitis models. Our study thus has unexpectedly identified cytosolic K6a as a novel negative regulator of TLR/NF-κB signaling preventing excessive proinflammatory cytokine and chemokine expressions. It further highlighted the functional importance of ELKS as a common signaling scaffold for both canonical and atypical IKK-dependent activation off NF-κB in corneal epithelial cells. These findings suggest that strengthening cytosolic K6a’s anti-inflammatory effect on epithelial TLR signaling could represent a potential therapeutic strategy for treating corneal inflammatory diseases, as well as for diseases where aberrant TLR activities have strong associations with disease pathogenesis and progression in other mucosal epithelia (Piccinini and Midwood 2010).

## Methods

### Cell culture and treatment

Telomerase-immortalized human corneal epithelial (Robertson et al. 2005) cells were grown in standard keratinocyte growth medium ^TM^ 2 (KGM^TM^-2) containing 0.15 mM CaCl_2_ (low Ca– KGM-2; Lonza) without gentamycin. 293T cells were maintained in Dulbecco’s Modified Eagle Medium containing 10% heat-inactivated fetal bovine serum (Corning). All cells were maintained in a humidified tissue culture incubator at 37°C and 5% CO_2_. *Pseudomonas aeruginosa* supernatant (Mc et al. 1999) was produced by inoculating the bacteria into tryptic soy broth (TSB) and incubated inside an orbital shaker at 180 rpm with aeration until the culture reached late log phase the next day. The bacteria were pelleted by centrifugation at 3,000 rcf for 20 min, followed by passing the supernatant through a 0.22μM polyether sulfone membrane filter (Millipore Sigma). KGM^TM^-2 containing 20% (v/v) of filtered PAO1 supernatant or TSB and KGM^TM^-2 containing purified flagellin at 1μg/ml or vehicle (tlrl-pafla; from *P. aeruginosa*; InvivoGen) were used for treating hTCEpi cells for various durations as indicated in the figure legends.

### Gene knockdown and CRISPR-knockout

For gene knockdown in hTCEpi cells, cells were seeded while at the same time being transfected with ON-TARGETplus non-targeting control (NTC) pool siRNAs (D-001810), K6a-, RelA-, ELKS-, or IKKε-specific SMARTpool siRNAs (L-012116; L-003533; L-010942; L-003723; Horizon Discovery) at 120nM each using Lipofectamine^TM^ RNAiMAX reagent (Invitrogen^TM^). Next day, the transfected cells were replenished with new KGM^TM^-2 and maintained for 2 more days for a total of 3 days before experiments. For CRISPR-genome editing studies, Alt-R^TM^ *S. pyogenes* Cas9 nuclease (version 3) was combined with duplexes of NTC- or K6a-specific Alt-R^TM^ CRISPR-Cas9 crRNAs pre-annealed with tracrRNAs (Integrated DNA technologies), forming Cas9-ribonucleic acid protein (RNP) complexes. Cas9-RNP complexes in solution R with 1.5 μM nuclease and 1.8 μM guide RNA were incubated for 30 mins before they were transfected into 3 million hTCEpi cells using the Neon transfection system and 10μl kit (Invitrogen^TM^). After transfection, the cells were seeded into a 10 cm^2^ tissue culture dish in KGM^TM^-2 and propagated for 1 week before the experiments.

### Cytokine antibody array and ELISA

hTCEpi cells transfected with siRNAs for 3 days were treated with KGM^TM^-2 containing 20% (v/v) TSB or sterile PAO1 supernatant for 24 hours. Conditioned media were harvested and the amount of IL-1α, IL-8 and CXCL1 cytokines within were quantified by ELISA according to manufacturer instructions (Duoset, R&D Systems). For cytokine antibody array, the conditioned media of NTC- and K6a-siRNA-transfected cells were mixed with a biotinylated detection antibody cocktail to form cytokine/ detection antibody complexes. After incubation with nitrocellulose membranes from the Human XL Cytokine Array kit (ARY022B, R&D Systems), the cytokine/ detection antibody complexes were bound by their cognate capturing antibodies immobilized on the membranes as spots. Following a wash to remove unbound material, streptavidin-horse radish peroxidases and chemiluminescent detection reagents were sequentially added to the membranes to generate light from these spots. As spot sizes were proportional to the amount of bound cytokine, they were imaged by the Syngene PXi Imager and quantified using the Image J software.

### RT-digital droplet PCR for mRNA transcript detection

To correlate proinflammatory cytokine transcript levels with the ELISA data, hTCEpi cells were stimulated by KGM^TM^-2 containing 20% (v/v) sterile PAO1 supernatant for 8 hours before lysis by TRI reagent (Ambion). Total RNAs were purified using Direct-Zol RNA miniprep kit (Zymo Research) and reverse-transcribed into cDNAs using cDNA iScript^TM^ advanced cDNA synthesis kit (Bio-Rad). Duplex ddPCR reactions were performed by combining cDNA products with human IL-1α, IL-8 or CXCL1 Taqman FAM-MGB probes together with TBP Taqman VIC-MGB probe (Hs00174092_m1, Hs00174103_m1, Hs00236937_m1, Hs00427620m1; Applied Biosystems) in 1X ddPCR^TM^ Supermix for probes (Bio-Rad) reaction mixture. Droplets were read by the QX200 platform and analyzed using the QX Manager program (Bio-Rad). Gene expression was normalized to TBP transcripts and expressed as fold change relative to NTC- and K6a-siRNA transfected cells under each treatment condition.

To evaluate the efficiency of mouse K6a knockout, K6a knockout and wildtype mice were sacrificed by cervical dislocation followed by eyeball enucleation. The corneas were excised out and the corneal epithelium were separated from the stroma by EDTA incubation at 37 °C for 1 hour (Spurr and Gipson 1985). Tissue lysis, total RNAs purification, reverse-transcription, singleplex ddPCR reactions using mouse K6a and GAPDH Taqman FAM-MGB probes (Mm00833464_g1, Mm99999915_g; Applied Biosystems), and droplet analysis were performed as above.

### Plasmids

Mammalian expression vectors employed in this study include pcDNA3.1-ELKS-Flag (Genscript), pcDNA3.1-IKKε-Flag (OHu2671; Genscript), pcDNA3.1-empty (Genscript), pEF1-empty-Myc-His (Invitrogen^TM^), pEF1-K6a (no tag), pCMV-HA_3_-K6a, pCMV-GFP-N1 (Clontech). pEF1-K6a (no tag) was constructed using a pUC57 vector encoding human K6a (codon-optimized by Genscript) as a PCR template to generate a full length human K6a DNA fragment terminated by a stop codon. The fragment was cloned into the pEF1-empty-Myc-His vector at the multiple cloning site upstream of the Myc-His tag sequence.

Lentiviral expression vectors employed in this study include pCDH533-Tet-On-IRES-Neo and pCDHTRE-K6a-mCherry. pCDH533-Tet-ON-IRES-Neo was constructed by cloning the reverse-tetracycline controlled transactivator from pTetOne (Clontech) into pCDH533-IRES-Neo (System Bioscience). For pCDHTRE-K6a-mCherry, it was constructed by first replacing the EF1a core/HTLV promoter from pCDH533-IRES-Neo with the TRE3GS promoter from pTetOne to create pCDHTRE-IRES-Neo. Second, a full length K6a sequence was cloned into pmCherry-N2 to form pmCherry-N2-K6a. Then, the entire K6a-mCherry sequence was synthesized by PCR from pmCherry-N2-K6a and cloned into pCDHTRE-IRES-Neo cut by Nhe1 and Sal1 to replace the IRES-Neo cassette.

### Lentiviral production

7 μg of lentiviral vectors was mixed with Lenti-X^TM^ packaging single shots (Clontech) in a final volume of 600μl for 15min before the transfection mix were added to 4 million of Lenti-X ^TM^ 293T cells seeded on a 10 cm^2^ tissue culture dish. The medium containing the virus particles was collected at 48h and 72 hours after transfection and pooled. After removal of cell debris by centrifugation at 500 rcf for 15 min, the clarified supernatant was combined with Lent-X^TM^ concentrator at a 3-to-1 volume ratio. After incubation for 1 hour, the lentivirus particles were pelleted by centrifugation at 1,200 rcf for 45 min at 4°C and resuspended in 500μL of KGM^TM^-2. The virus titer was determined using Lenti-X^TM^ GoStix Plus (Clontech).

### Inducible expression of K6a-mCherry in hTCEpi cells

0.1 million of hTCEpi cells seeded on 24-well plate were transduced by VSV-G lentiviral particles encoding pCDH533-Tet-On-IRES-Neo at 40 multiplicities of infection in the presence of 2 μg/ml polybrene in KGM^TM^-2. Transduction was terminated after two hours by washing the cells two times with KGM^TM^-2. Cells were expanded and propagated for 1 week before G418 selection. G418-resistant cell pool were transduced by VSV-G lentiviral particles encoding pCDHTRE-K6a-mCherry as above and propagated without selection. To induce K6a-mCherry expression, cells were treated with 1 μg/ml doxycycline for 24 hours.

### Western blot and immunoprecipitation antibodies

Rabbit polyclonal antiserums raised against the 19-mer KAMP peptide (RAIGGGLSSVGGGS STIKY; New England Peptide) or the 10-mer KAMP peptide (GGLSSVGGGS; Pacific Immunology), which correspond to K6a at the carboxyl terminal region between amino acid residues 533–551 and 537–546 respectively, have been described previously (Chan et al. 2018). Both rabbit polyclonal antiserums recognize human K6a. The rabbit anti-10-mer serum was further purified using Protein A IgG purification kit and conjugated to agarose beads by AminoLink^TM^ Immobilization kit (Thermo Scientific ^TM^ Pierce ^TM^). Other rabbit polyclonal antibodies included anti-ELKS and anti-IKKε antibodies (clone D20G4, clone P85; Cell Signaling Technology). Rabbit monoclonal antibodies included anti-PO4 IKKα/β Ser-176/ Ser-180 and anti-PO4 IKKε Ser-172 antibodies (clone 16A6, clone D1B7; Cell Signaling Technology). Mouse monoclonal antibodies included anti-K6a antibody (clone B7; Santa Cruz Biotechnology), anti-RelA antibody (clone A21012A; BioLegend), anti-PO4 IκBα Ser-32/Ser-36 and anti-β-actin antibodies (clone 5A5, clone 3700; Cell Signaling Technology), THE^TM^ DYKDDDDK tag antibody (A00188; Genscript) and Direct-Blot^TM^ HRP anti-mCherry antibody (clone 8C5.5; BioLegend).

### Liquid chromatography–tandem mass spectrometry (LC-MS) analysis

hTCEpi multi-layered cultures (Chan et al. 2018) grown in a 3-μM polyester 6 well transwell plate (Corning Costar) were harvested and lysed by cell lysis buffer consisted of 1% (v/v) NP-40 in 20 mM Hepes, pH 7.2, 120 mM NaCl, 1 mM EDTA, and 1× Halt^TM^ protease and phosphatase inhibitor cocktail (Thermo Scientific ^TM^) for 15 min on ice. The lysate was clarified by centrifugation at 15,000 rcf for 30 min at 4°C and the supernatant was mixed with 30 µL of agarose beads conjugated to purified anti–10-mer KAMP antibodies overnight 4°C with rotation. After removal of unbound material and four washes with lysis buffer, bound proteins were eluted from the beads by incubation with free 10-mer KAMP peptides at 1 μg/ml for 15 min at room temperature. The eluate was resolved in a 12% Bis-Tris protein gel (NuPAGE), followed by gel-staining by Coomassie Blue dye. Each gel band of interest was excised and digested with trypsin or chymotrypsin for peptide extraction. Extracted peptides were resuspended in 30μl in 1% acetic acid and injected into an Acclaim ^TM^ PepMap ^TM^ 100 Å–pore-size C18 reversed phase capillary chromatography column (Thermo Scientific ^TM^) programmed for running in an acetonitrile/ formic acid gradient at a 0.25μl/min flow rate. Peptides eluted from the column were introduced into a Finnigan LTQ-Obitrap Elite hybrid mass spectrometer (Thermo Scientific ^TM^) with electrospray ion source setting at 2.5 kV. Collision-induced dissociation spectra were collected and searched against the human reference sequence database using the Mascot program and more specifically against the human K6a sequence using the Sequest program.

### Western Blot analysis

After stimulation with PAO1 supernatant or flagellin, hTCEpi cells were washed with iced cold PBS, and lysed in buffer consisted of in 50 mM Tris, pH 7.2, 150 mM NaCl, 1 mM EDTA, 1% (v/v) Tx-100 and 1× Halt^TM^ protease and phosphatase inhibitor cocktail for 30 min on ice. Lysate samples were clarified by centrifugation at 15,000 rcf for 30 min at 4°C and their protein concentration was determined by BCA assay (Thermo Scientific ^TM^ Pierce ^TM^). Samples normalized to equal amounts were denatured in Laemmli’s buffer at 95°C for 10 min and resolved by SDS-PAGE using Criterion ^TM^ TGX ^TM^ Midi protein gels (Bio-Rad). Resolved proteins were transferred onto 0.2μM PVDF membrane by the Criterion ^TM^ Blotter (Bio-Rad). Proteins of interest were immunoblotted using their respective antibodies or antiserum and detected by chemiluminescence using Pierce ^TM^ ECL Western blotting (Thermo Scientific ^TM^) or Trident femto Western HRP (GeneTex) substate.

### Immunoprecipitation

To assess the interaction between endogenous K6a and ELKS, 1 mg of hTCEpi cell lysate in cell lysis buffer was precleared with 25 μl of protein A/G plus agarose (SC-2003; Santa Cruz Biotechnology) for 2 hours at 4°C with rotation before the agarose was removed by centrifugation at 2,000 rcf for 10 min. The precleared lysate was retrieved and mixed with 20 mg of monoclonal anti-K6a antibodies overnight followed by incubation with 30 μl of protein A/G magnetic beads (Thermo Scientific ^TM^ Pierce ^TM^) for 1 hour. After four washes in cell lysis buffer, immunoprecipitated proteins were eluted from the beads by incubating in 60 µl of Laemmli’s buffer at 95°C for 10 min. Eluted samples were resolved in Mini-Protean TGX 7.5% precast gel (Bio-Rad) and immunoblotted with rabbit polyclonal anti-ELKS antibody and anti– 19-mer KAMP antiserum for the detection of ELKS and K6a respectively.

To assess the interaction between overexpressed K6a and ELKS, 293T cells were transfected with pcDNA3.1-ELKS-Flag (Genscript), pEF1-K6a (no tag), pCMV-HA_3_-K6a and empty vectors as indicated in figures 3A and B for 24 hours followed by cell lysis. K6a and ELKS were immunoprecipitated from precleared lysate using anti–19-mer KAMP antiserum and monoclonal THE^TM^ DYKDDDDK tag antibody respectively followed by immunoblotting for the co-immunoprecipitated ELKS and K6a using THE^TM^ DYKDDDDK tag antibody and anti–19-mer KAMP antiserum. To evaluate the interaction between the overexpressed ELKS and IKKε, 293T cells were transfected with pcDNA3.1-ELKS-Flag, pcDNA3.1-empty, pcDNA3.1-IKKε-Flag and pcDNA3.1-empty as indicated in figures 5C and D for 24 hours followed by cell lysis. ELKS and IKKε were immunoprecipitated from the precleared lysate using rabbit polyclonal anti-ELKS and anti-IKKε antibodies respectively followed by immunoblotting for the co-immunoprecipitated ELKS and IKKε using polyclonal anti–IKKε and anti-ELKS antibodies.

### Generation of corneal epithelium-specific and whole body K6a knockout mice

To create floxed-K6a mice, embryonic stem (C57BL/6) cells were first electroporated with a linearized vector which encodes the 5’ and 3’ homology arms with lox P sites flanking the first and last K6a exons and a neomycin-resistant gene cassette that is flanked by flippase (Flp) recognition target sites and positioned upstream of the 3’ lox P site (Cyagen). After neomycin selection, 45 drug-resistant clones were derived, 2 clones were confirmed as being correctly targeted by Southern Blotting, and 1 clone (1B3) was injected into host embryos for chimera production in a surrogate mother. Male founder chimeras were bred to wildtype female Flp-deleter mice to remove the neomycin-resistant gene and yield F1 mice that were heterozygous for the floxed-K6a allele (3 males and 4 females), which were then bred to generate progenies homozygous for floxed-K6a (K6a ^F/F^).

To generate corneal epithelium-specific K6a knockout mice, K6a ^F/F^ mice were first crossed with C57BL/6 driver mice that express the Cre recombinase under the keratin 12 promoter (K12-Cre ^Tg/null^; RRID: IMSR_Jax: 023055, The Jackson Laboratory). This promoter was originally chosen because K12 is widely accepted as the differentiation marker in adult corneal epithelium (Hayashi et al. 2010; Kao 2020), but it was found leaky within the gametes of Lac-EGFP reporter mice with higher penetrance in male (46%) than in female (7%) (Weng et al. 2008). As such, we adopted a recommended strategy where only female mice carrying a single copy of K12-Cre (K12-CRE ^Tg/null^) were used as breeders (Weng et al. 2008; Song and Palmiter 2018).

After the first cross, mice heterozygous for floxed-K6a (K12-Cre ^Tg/null^; K6a ^F/WT^) were bred with K6a ^F/F^ mice again to obtain the desired homozygous floxed-K6a mice (K12-Cre ^Tg/null^; K6a ^F/F^). Because some of the progenies still developed ectopic K6a deletion(s) as revealed by ear tissue genotyping (i.e. K6a ^F/F^ becomes K6a ^D/D^ outside the corneal epithelium), suggesting Cre activity during embryogenesis, we ensured that when compared between corneal epithelium-specific K6a knockout mice (K12-Cre ^Tg/null^; K6a ^F/F^) and floxed-K6a control mice (K12-Cre ^null/null^; K6a ^F/F^), all ectopic K6a-deleted mice (K6a ^D/D^) were being identified by detailed genotyping analysis and excluded in the experiment.

To generate whole body K6a knockout mice, K12-Cre ^Tg/null^; K6a ^D/D^ mice were backcrossed to C57BL/6 mice (The Jackson Laboratory) for several generations to remove the K12-Cre transgene but retain the K6a ^D^ allele followed by inbreeding to generate mice with homozygous K6a knockout (K12-Cre ^null/null^, K6a ^D/D^; or K6a ^D/D^ for short). Reverse transcription-ddPCR was employed to confirm that the corneal epithelium of K6a knockout mice did not express K6a mRNA transcripts (Supplementary Figure 1A).

### Genotyping

The following primers were used to distinguish among K6a ^WT^, K6a ^F^ and K6a ^D^ alleles: K6a region 1 forward primer (5’ loxP site 1): 5’-*ACA CAG GGA GTG TCT ACA TGG GCA*-3’^;^ K6a region 1 reverse primer (3’ loxP site 1): 5’-*GCA AAT TCT CCT GCC TTC CAG C*-3’; K6a region 2 forward primer (5’ loxP site 2): 5’-*CAG ATG TCA GTT CAG CAT TCC TTC*-3’; K6a region 2 reverse primer (3’ loxP site 2): 5’-*ATT GTG GCT TCT TGT CAA GCA A*-3’; K6a deleted forward primer (5’ K6A region 1 forward primer): 5’-*ATG TAT GGC TTC AGG GAG AGA CTT AG*-3’^;^ K6a deleted reverse primer (3’ K6A region 2 reverse primer): 5’-*TGT TGT GTT TCT GCT GTC CCT TTA G*-3’. PCR reactions were performed using the Phire tissue direct PCR master mix (Invitrogen^TM^) with the program set at 98 °C for 5 min followed by 40 cycles of 98 °C for 5 sec, 63.2 °C for 5 sec and 72 °C for 20sec, followed by final extension at 72 °C for 10min. The following primers were used for tracking the insertion or elimination of the IRES-Cre transgene at the 3’ untranslated region of mouse K12 genomic locus: K12-Cre ^null^ forward primer (intron 7): 5’-*TTT GGA ATG GAG TCA CTT GC*-3’; K12-Cre ^Tg^ forward primer (Cre ORF): 5’-*GCA ACA GAG TTA GGA CTT GAA CCC*-3’; K12-Cre ^Tg/null^ common reverse primer (3’UTR): 5’-*AAA GCG CAT GCT CCA GAC TGC C*-3’. PCR reactions followed the protocol posted at: https://www.jax.org/Protocol?stockNumber= 023055&protocolID=25473. Expected results were shown in Supplementary Figure 1B.

### Murine models of corneal inflammation and infection

All procedures performed were in compliance with the Public Health Service Policy on Humane Care and Use of Laboratory Animals, and were approved by the Institutional Animal Care and Use Committee of the Cleveland Clinic. Both female and male mice (10-12 weeks old) were used. Anesthesia was induced by intraperitoneal injection (50 μl per 25 g body weight) of ketamine [50 mg/kg body weight (BW)] and dexmedetomidine (0.375 mg/kg BW) cocktail and reversed by atipamezole (3.75 mg/kg BW). The epithelial barrier of the mouse cornea was compromised by three parallel abrasions with a 26-gauge needle before inoculation. In the sterile inflammation model, purified *P. aeruginosa* LPS or *S. aureus* LTA (20 μg in 2 μl PBS; Sigma-Aldrich) was topically inoculated to the scarified corneas of C57BL6 wildtype mice (K6a ^WT/WT^) and whole body K6a knockout mice (K6a ^D/D^) for 3 or 24 hours. In the bacterial infection model, live *S. aureus* (10^6^ CFU in 5 μl saline) or *P. aeruginosa* PA3346 (10^5^ CFU in 5 μl saline) was inoculated onto the scarified corneas of corneal epithelium-specific K6a knockout mice (K12-Cre ^Tg/null^; K6a ^F/F^) and floxed-K6a mice (K12-Cre ^null/null^; K6a ^F/F^) as controls (Song and Palmiter 2018). The disease progression in infected corneas was examined at 24 hours under a stereomicroscope equipped with digital camera. A five-point grading system (0 to 4 points) was used to assess the disease severity based on the area of opacity, density of central opacity, density of peripheral opacity and surface irregularity in a blind-folded manner followed by summation for a total disease score (0-16 points).

At the end of experiments, mice were euthanized immediately before their whole corneas were collected by dissection and homogenized in sterile PBS (150 μl each) using Precellys CK14 ceramic beads and tissue homogenizer (Bertin Instruments). After removal of debris by centrifugation, the cytokines within the homogenate were measured by ELISA (DuoSet, R&D Systems) using half area 96-well microplate (Corning Costar). Viable bacterial loads were quantified by serial dilution and plating on tryptic soy agar.

### Flow cytometric analysis of neutrophils and macrophages

For flow cytometric analysis of immune cells, mouse corneas were incubated in type I collagenase (Sigma Aldrich) at 82 U/ cornea for 2 hours at 37 °C, followed by passing the cell suspension through a 30 μm filter to remove any residual undigested tissues. Cells were then washed and incubated at 4 °C for 10 min in 100 μl ice cold FACS buffer (PBS with 1% FBS) containing 2 μg of Fc blocker (anti-mouse CD16/CD32 antibody; eBioscience). Antibody staining was performed on ice for 1 hour with APC/Cy7 anti-mouse CD45 (clone 30-F11; BioLegend), rat monoclonal FITC anti-mouse Ly6B.2 (clone 7/4; Abcam) and PE anti-mouse F4/80 (clone BM8; BioLegend). All samples were stained with Zombie Violet (BioLegend) to gate out dead cells. Stained cells were washed twice with FACS buffer and resuspended in 0.5% PFA for detection with a BD Fortessa flow cytometer. Cells were first gated using forward scatter (FSC) and side scatter (SSC), then gated on viability, CD45+ leukocyte population, and cell type markers (Ly6B.2 and F4/80). Unstained samples and fluorescence-minus-one controls were used to confirm the gating strategy. Analysis of flow cytometric data was performed using the BD FlowJo software (version 10).

### Statistics

GraphPad Prism (version 9) was used to perform statistical tests. Normal distribution of data was determined by Shapiro-Wilk test. Equality of variances was determined by Brown-Forsythe test. Outliers were detected by inter-quantile range test. For animal studies, two groups of data with normal distribution were compared by unpaired t test for equal variances and by Welch’s t test for unequal variances. For *in vitro* cell culture studies, data groups with normal distribution were compared by one-way ANOVA followed by Šidák’s post hoc comparisons. A significance threshold of P < 0.05 was considered statistically significant.

## Results

### K6a knockout corneas substantially produce LPS-induced cytokines and chemokines in the presence of anti-inflammatory KAMPS

We first evaluated the effects of K6a knockout to cytokine and chemokine production by the corneal epithelium in an established mouse model of sterile inflammation (Sun et al. 2023). We compared scarified corneas inoculated by purified bacterial ligands (LPS) for 3 hours between whole body K6a knockout (K6a ^D/D^) mice and wild type (K6a ^WT/WT^) mice. We chose for a short inoculation time because the main focus was to assess the early productions of pro-inflammatory cytokines and chemokines by the corneal epithelium before myeloid immune cell infiltration from the surroundings (de Paiva, St Leger, and Caspi 2022). We isolated whole corneas from euthanized mice and immediately quantified their cytokines by ELISA. In response to LPS, the corneas of K6a knockout mice produced more cytokines (IL-6, GM-SF, TNFα) and chemokines (CXCL1, CXCL2, CXCL10) than those from wild type mice (Figure 1; left vs right black bars). Consistent with our previous report on KAMP’s anti-inflammatory effect on myeloid immune cells (Sun et al. 2023), the corneas of wild type mice strongly reduced their cytokine expressions if they were pre-treated topically with a high dose of KAMP10 (K10) before LPS inoculation (Figure 1; left magenta bars); an effect not seen with the scramble KAMP10 (SC10; Figure 1; left grey bars). Surprisingly, while KAMP10 was also suppressive in K6a knockout corneas, these corneas had retained considerable capacity for cytokine and chemokine productions that were comparable to corneas from wildtype mice (Figure 1; left black vs right magenta bars). These results therefore argue that beyond the anti-inflammatory and antimicrobial activities of KAMPS, the cytosolic K6a within the corneal epithelium itself provides additional anti-inflammatory function *in vivo*. In addition, unlike the non-K6a expressing myeloid immune cells which are only responsive to the anti-inflammatory effect of KAMPS, the corneal epithelium is responsive to both KAMPS- and cytosolic K6a-mediated suppression of pro-inflammatory cytokine and chemokine productions (Figure 1; left vs right magenta bars).

**Figure 1.**
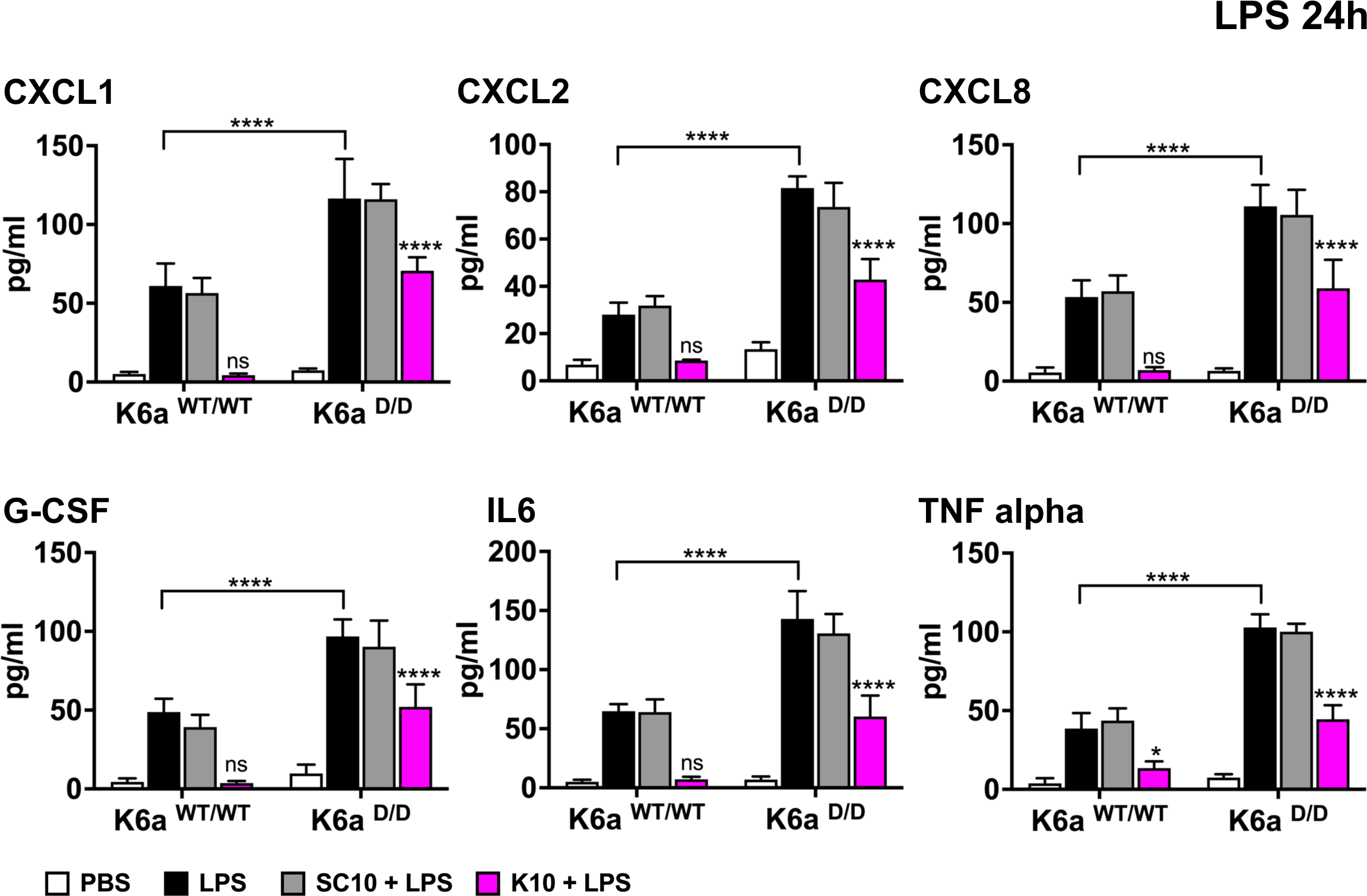
K6a knockout corneas substantially produce LPS-induced cytokines and chemokines in the presence of anti-inflammatory KAMP. Scarified mouse corneas (one per mouse) of C57BL/6 wildtype (K6 ^WT/WT^) mice and whole body K6a knockout (K6a ^D/D^) mice were sham-treated (black bars) or given one topical application of scramble KAMP10 peptide (SC10; grey bars), or KAMP10 peptide (K10; magenta bars) at 100 μg/ml for 30 min before inoculation with purified LPS (20 μg; black, grey, magenta bars) or PBS (white bar) for 3 hours. The KAMP dosage was chosen based on previous *in vitro* data that effectively abolished LPS-induced inflammatory response in murine neutrophils and macrophages (Yan et al., 2023). Mice were then euthanized and whole corneas were dissected for cytokine and chemokine quantification by ELISA. Means ± SD (n = 7 to 9 mice (corneas) per group). *P < 0.05, **P < 0.01, ***P < 0.001, and ****P < 0.0001. ns, P > 0.05. Statistical significance was determined by one-way ANOVA followed by Šidák’s post hoc comparison to PBS inoculation within the same group (asterisks without bracket) or between LPS-inoculated wildtype and K6a knockout mice (asterisks with brackets).

### K6a suppresses NF-κB/ RelA-dependent expressions of proinflammatory cytokines in corneal epithelial cells

To discern the molecular underpinning of cytosolic K6a’s anti-inflammatory action at the corneal epithelium, we hypothesized that it involves antagonism towards the activity of NF-κB, since this transcription factor family is well-known for driving corneal inflammation in response to TLR activation (Pearlman et al. 2013; Fortingo et al. 2022). We focused on three NF-κB targets: IL-1α, IL-8, CXCL1, based on preliminary results that these pro-inflammatory cytokines were upregulated in human telomerase-immortalized corneal epithelial (hTCEpi) cells with K6a knockdown (KD) as assayed by cytokine antibody array (Supplementary Figure 2). We assessed whether K6a suppresses their productions by specifically inhibiting RelA; the prototypical NF-κB member activated after TLR stimulation in the corneal epithelium (Kumar et al. 2007; Zhang et al. 2013; Wu et al. 2021). We transfected non-targeting control (NTC) siRNAs or K6a-specific siRNAs alone, or together with RelA-specific siRNAs into hTCEpi cells for 3 days, followed by cell treatment with media containing 20% (v/v) bacterial tryptic soy broth (TSB) as a basal uninduced control or 20% (v/v) sterile supernatant from *Pseudomonas aeruginosa* culture harvested at late log-phase (PAO1) for combined bacterial ligand TLR stimulations (Ino et al. 1999; Mc et al. 1999; Hayashi et al. 2001; Simon et al. 2014). We collected the conditioned KGM2 media and performed ELISA to quantify the cytokines secreted by the cells (Figure 1A).

In addition, we extracted their RNA and measured the cytokine mRNA transcript expression (Figure 1B). Under both basal and PAO1 stimulated conditions in the K6a KD cells, more IL-1α, IL-8 and CXCL1 cytokines (Figure 1A; blue vs magenta bars) and their transcripts were detected (Figure 1C; blue vs magenta bars). In contrast, K6a- and RelA double KD cells markedly reduced pro-inflammatory cytokine secretions and mRNA expressions compared to cells with K6a KD alone. (Figures 1A and 1B; magenta vs cyan bars). These findings therefore revealed that under both conditions the expression of these pro-inflammatory cytokines was driven by RelA but suppressed by K6a, supporting the idea that K6a antagonizes the TLR signaling pathway that leads to the activation of RelA activity.

### K6a interacts with canonical IKKα/β regulator ELKS to downregulate proinflammatory cytokine and chemokine expressions

We next performed a proteomic screen using hTCEpi cells to identify K6a-interacting proteins that functionally regulate NF-κB. We immunoprecipitated K6a present in whole cell lysates from uninduced cells followed by mass spectrometry analysis to identify co-eluted factors. We found in addition to well-known K6a-interacting partners such as K16, K17 and 14-3-3 proteins, several factors related to the NF-κB signaling pathway were present (Table 1). These include ELKS/Rab6-interacting/CAST family member 1 (ELKS), carboxyl terminus of constitutive heat shock cognate 70 (HSC70)-interacting protein (CHIP), tripartite motif-containing protein 29 (TRIM29), and RelA-associated inhibitor. ELKS forms a stable complex with NEMO; an essential adaptor where canonical IKKα/β associate and become activated. Functionally, ELKS bridges IκBα with NEMO enabling IKKα/β to phosphorylate IκBα at serine residues 32 and 36 (Ducut Sigala et al. 2004). Phosphorylated IκBα then becomes ubiquitinated and degraded, which releases RelA for transactivation activity in the nucleus. Likewise, CHIP and TRIM29 are E3 ligases with opposing actions onto NF-κB activity. While CHIP promotes NF-κB activation in dendritic cells by K63-linked polyubiquitination of TLR4/9 cofactors Src and PKCζ kinases (Yang et al. 2011), TRIM29 represses TLR4 signaling in alveolar macrophages by K48-linked polyubiquitination of NEMO leading to its degradation (Xing et al. 2016). RelA-associated inhibitor suppresses the DNA binding activity of RelA (Yang et al. 1999; Takada et al. 2002).

**Table 1.**

| Protein | kDa | Uniprot # | # of Peptides | Sequence coverage | Mascot score |
| --- | --- | --- | --- | --- | --- |
| Keratin, type II cytoskeletal 6A | 60 | P02538 | 42 | 52% | 1.78 |
| Keratin, type I cytoskeletal 16 | 51 | P08779 | 16 | 59% | 0.569 |
| Keratin, type I cytoskeletal 17 | 48 | Q04695 | 14 | 51% | 0.556 |
| 14-3-3 protein sigma | 28 | P31947 | 14 | 72% | 0.374 |
| 14-3-3 protein beta/alpha | 28 | P31946 | 6 | 54% | 0.0559 |
| 14-3-3 protein theta | 28 | P27348 | 7 | 53% | 0.102 |
| 14-3-3 protein gamma | 28 | P61981 | 6 | 42% | 0.0487 |
| 14-3-3 protein zeta/delta isoform X1 | 42 | P29310 | 11 | 35% | 0.0851 |
| ELKS/Rab6-interacting/CAST family member 1 isoform epsilon | 128 | Q8IUD2 | 36 | 27% | 0.214 |
| E3 ubiquitin-protein ligase CHIP | 35 | Q9UNE7 | 4 | 13% | 0.077 |
| Tripartite motif-containing protein 29 isoform X1 | 66 | Q14134 | 7 | 12% | 0.036 |
| RelA-associated inhibitor isoform X1 | 96 | Q8WUF5 | 4 | 6.20% | 0.008 |

We decided to focus on ELKS because it has the highest peptide spectral coverage according to the mass spectrometry results (Table 1). We overexpressed K6a and ELKS proteins in 293T cells and confirmed their interaction in forward and reverse co-immunoprecipitation (Co-IP) experiments (Figures 3A and 3B). We further validated their interactions by Co-IP studies using lysates from hTCEpi cells containing stable expression of mCherry-K6a or endogenous K6a (Figures 3C and 3D). Furthermore, double KD of K6a and ELKS in hTCEpi cells, similar to the double KD of K6a and RelA, significantly suppressed the increased pro-inflammatory cytokine secretion observed in cells with K6a KD alone under both basal and PAO1-stimulated conditions (Figures 2E, magenta vs cyan bars). Together, these results support the notion that K6a exerts its negative regulation of NF-κB by interacting with ELKS.

**Figure 2.**
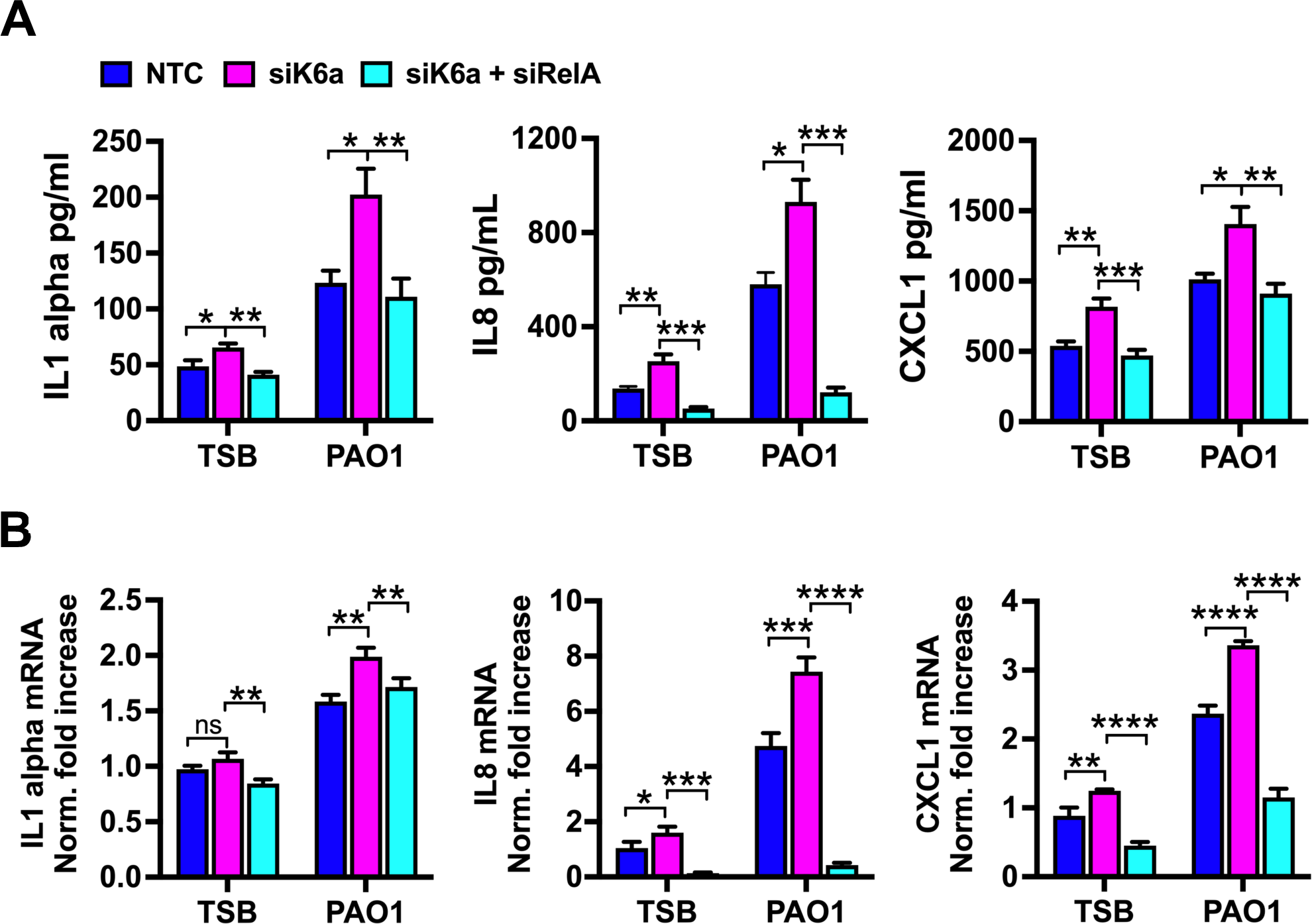
K6a inhibits NF-κB/ RelA-dependent cytokine expressions in corneal epithelial cells. **(A and B)** hTCEpi cells transfected with non-targeting control (NTC), K6a-, and RelA-specific SMARTpool siRNAs as indicated for 3 days were treated with KGM^TM^-2 tissue culture media containing 20% (v/v) bacterial tryptic soy broth (TSB) as basal uninduced control or 20% (v/v) sterile *Pseudomonas aeruginosa* (PAO1) supernatant for mixed bacterial ligand stimulations for (A) 24 hours and 8 hours (B) respectively. (A) Conditioned KGM^TM^-2 media were collected for ELISA quantifications of cytokines. (B) Total RNA was purified from cells, reverse-transcribed and assayed for cytokine mRNA transcript expressions by digital droplet PCR. Gene expression was normalized to TBP transcripts and expressed as fold change relative to NTC- or K6a-siRNA transfected cells under each treatment condition. Means ± SEM (n > 3) for A and ± SD (n = 3) for B. *P < 0.05, **P < 0.01, ***P < 0.001, and ****P < 0.0001. ns, P > 0.05. Statistical significance was determined by one-way ANOVA followed by Šidák’s post hoc pairwise comparison to NTC or K6a-siRNA transfected cells within each treatment group.

**Figure 3.**
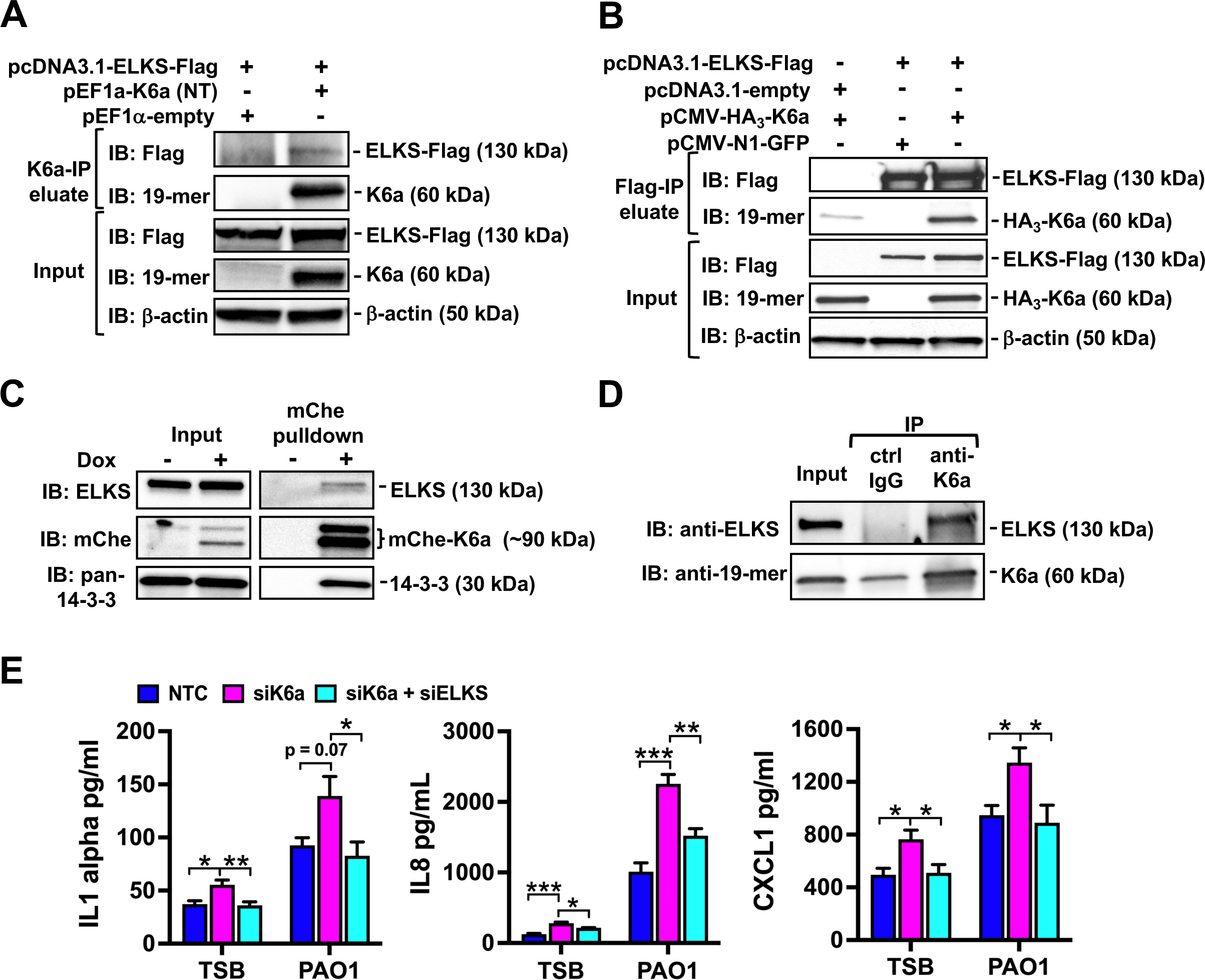
K6a interacts with canonical IKKα/β regulator ELKS to attenuate pro-inflammatory cytokine expressions. **(A and B)** 293T cell lysates containing overexpressed Flag-tagged ELKS and K6a in combinations indicated were immunoprecipitated (IP) with mouse monoclonal anti-K6a antibody (A) and THE^TM^ DYKDDDDK tag mouse monoclonal antibody (B) respectively. Eluted samples were immunoblotted for the detection of ELKS using THE^TM^ DYKDDDDK tag antibody (A and B) and K6a using rabbit polyclonal anti-19-mer serum (B) respectively. **(C)** hTCEpi cells stably expressing Tet-On transactivator and K6a-mcherry driven by Tet-On responsive promoter were sham- or doxycycline-induced at 1 μg/ml for 24 hours followed by cell lysis. K6a-mCherry fusion protein in lysates was pulled down by nanobodies conjugated to Chromotek RFP-Trap agarose. Eluted samples were immunoblotted with rabbit polyclonal anti-ELKS and Direct-Blot^TM^ HRP anti-mCherry antibodies for the detection of ELKS and K6a-mCherry respectively. **(D)** Endogenous K6a in hTCEpi cell lysates was immunoprecipitated using mouse IgG control and monoclonal anti-K6a antibody. Eluted samples were immunoblotted with rabbit polyclonal anti-ELKS antibody and anti-19mer serum for the detection of ELKS and K6a respectively. **(E)** hTCEpi cells transfected NTC-, K6a-, and ELKS-specific SMARTpool siRNAs as indicated for 3 days were treated with KGM^TM^-2 media containing 20% (v/v) TSB or sterile PAO1 supernatant for 24 hours, followed by conditioned media collection for ELISA. Means ± SEM (n > 3). *P < 0.05, **P < 0.01, ***P < 0.001, and ****P < 0.0001. ns, P > 0.05. Statistical significance was determined by one-way ANOVA followed by Šidák’s post hoc pairwise comparison to NTC or K6a-siRNA transfected cells within each treatment group.

### K6a does not inhibit canonical IKKα/β-dependent phosphorylation and degradation of IκBα

As mentioned above, IKKα/β phosphorylates IκBα to promote its degradation and the release of NF-κB into the nucleus to induce transcription. The immediate target of NF-κB is IκBα itself, which associates and sequesters NF-κB back into the cytoplasm terminating the transcriptional response. During sustained stimulation, these events repeat as NF-κB oscillations and eventually dampen out depending on the stimulus strength (DeFelice et al. 2019; Kizilirmak, Bianchi, and Zambrano 2022). Intermediate stimulus promotes multiple rounds of steadfast NF-κB activities while strong stimulus promotes NF-κB activities with high initial amplitude but dampens out shortly (DeFelice et al. 2019). We discovered that purified TLR5 agonist flagellin was a strong stimulus while the non-purified mixed TLR agonists from PAO1 supernatant was intermediate. Specifically, NTC-control hTCEpi cells treated with flagellin underwent one round of robust phosphorylation of IKKα/β at 20 min (Figure 4A, left panel, lane 2, column 2) preceding peak IκBα phosphorylation at 1 hour (Figure 4A, left panel, lane 3, column 3). These events resulted in full IκBα degradation as early as 20 min followed by its complete resynthesis at 2 hours (Figure 4A, left panel, lane 3, columns 2 and 4), indicating one cycle of robust NF-κB activity. In contrast, NTC-control hTCEpi cells stimulated with PAO1 underwent two rounds of IκBα phosphorylation at 1 hour and 6 hours respectively (Figure 4B, left panel, lane 2, columns 3 and 5) and were accompanied with subtle but noticeable IκBα degradation and resynthesis at 1 hour and 3 hours respectively (Figure 4A, left panel, lane 3, columns 3 and 4), suggesting that NF-κB activity had cycled twice during the time intervals examined.

**Figure 4.**
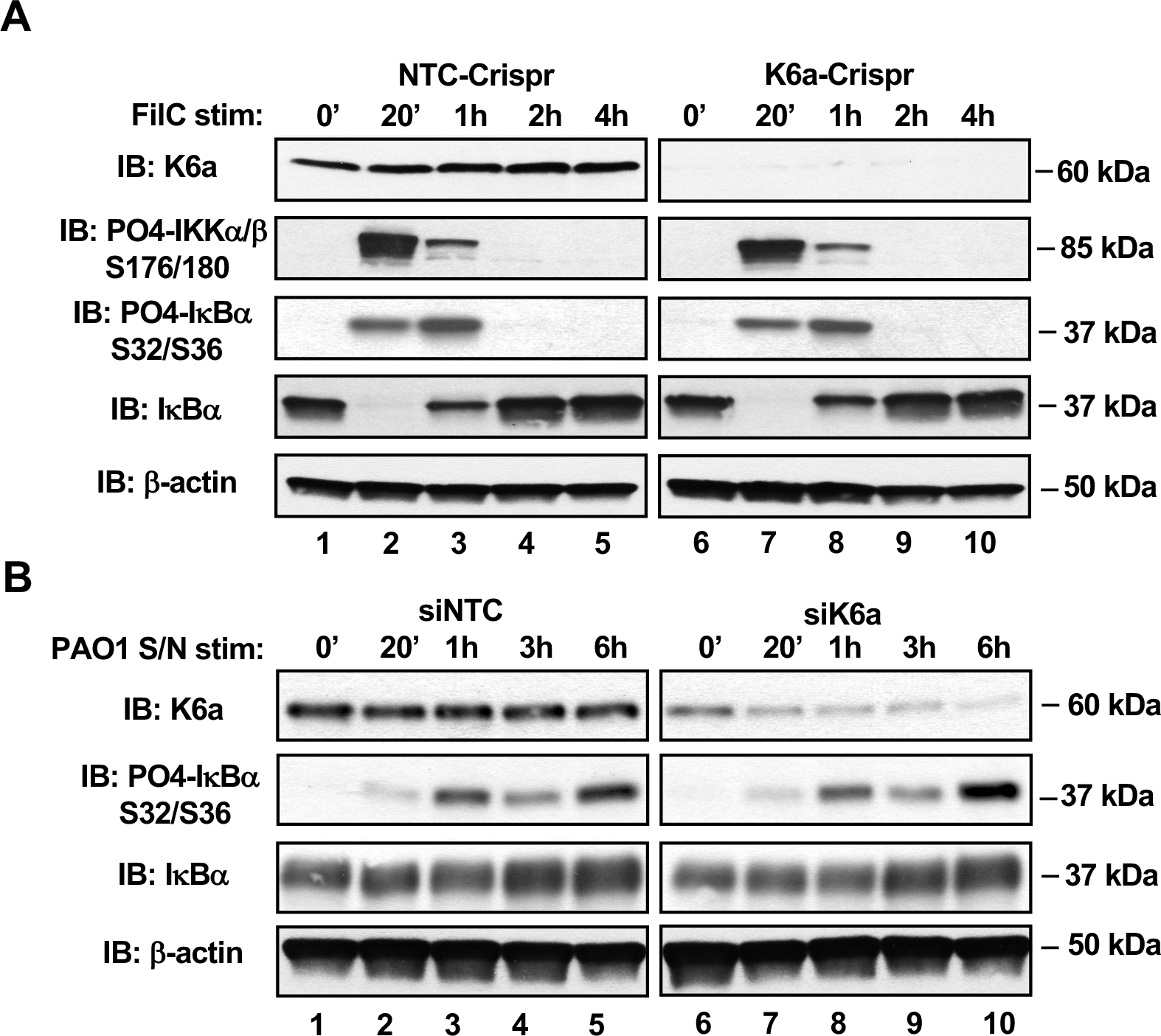
K6a does not inhibit canonical IKKα/β-dependent phosphorylation and degradation of IκBα. **(A)** hTCEpi cells introduced with CRISPR-Cas9 NTC- or K6a-specific guide RNA by Neon transfection for 1 week were treated with purified 500 ng/ml PAO1 flagellin for 0 min, 20 min, 1 hour, 2 hours, and 4 hours before cell lysis. Equal amounts of cell lysate at each time point were immunoblotted for PO4-IKKα/β Ser-176/ Ser-180, PO4-IκBα Ser-32/Ser-36, total IκBα, K6a and β-actin by their respective antibodies. (**B)** hTCEpi cells transfected with NTC- or K6a-siRNAs for 3 days were treated with KGM^TM^-2 media containing 20% (v/v) sterile PAO1 supernatant for 0 min, 20 min, 1 hour, 3 hours, and 6 hours before cell lysis. Equal amounts of cell lysate at each time point were immunoblotted for the detection of PO4-IκBα Ser-32/Ser-36, total IκBα, K6a and β-actin by their respective antibodies.

We next compared the kinetics of IκBα phosphorylation and protein levels between NTC control and K6a KD hTCEpi cells during flagellin or PAO1 stimulation. We hypothesized that the increased pro-inflammatory cytokine expressions we observed in K6a KD cells under the basal condition was due to autonomous activation of the canonical NF-κB signaling pathway when K6a was knocked down. As such, we would expect to find a higher level of phosphorylated IκBα and a lower steady-state level of IκBα within these cells in the absence of stimulation compared to NTC control cells. Unexpectedly, we found that in both cell populations IκBα had remained unphosphorylated at time 0 (Figures 4A and 4B, lanes 3 and 2, columns 1 vs 6) and its initial levels were the same (Figures 4A and 4B, lanes 4 and 3, columns 1 vs 6). Moreover, after flagellin or PAO1 stimulation, both the kinetics of IKKα/β phosphorylation (Figure 4A, all lane 2), IκBα phosphorylation (Figures 4A and B, all lanes 3 and 2) as well as IκBα degradation and resynthesis (Figure 4A and B, all lanes 4 and 3) were comparable between NTC control versus K6a KD cells. These results therefore suggest that the interaction between K6a and ELKS is unlikely to interfere with the activating effects of ELKS on the canonical IKKα/β-dependent phosphorylation of IκBα and its downstream signaling events.

### K6a suppresses ELKS-associated atypical IKKε in autophosphorylation and phosphorylation of RelA

We then evaluated how K6a antagonizes NF-κB activation by exploring ELKS-regulated pathways that are independent of IκBα phosphorylation and degradation. Notably, the IKK family in addition to canonical IKKα/β include the non-canonical IKK members IKKε and TBK1. After TLR stimulation in embryonic fibroblasts and macrophages, TBK and IKKε are critical for inducing type-I interferon response but they are dispensable for NF-κB activation (Hemmi et al. 2004; Perry et al. 2004). In contrast, IKKε-induced NF-κB activity is crucial for viral-G protein coupled receptor signaling in tumorigenesis and in the nuclear retention of NF-κB in activated T cells (Peters, Liao, and Maniatis 2000; Mattioli et al. 2006; Wang et al. 2013). Moreover, RelA activation in a number of cell types has been reported to bypass IκBα degradation where RelA appears to be more abundant than IκBα and contains a sub-pool free from IκBα sequestration (Buss et al. 2004; Douillette et al. 2006; Choudhary et al. 2007; Sitcheran et al. 2008). How RelA is maintained transcriptionally latent in such sub-pool is not fully understood but one mechanism could involve the binding of nuclear long non-coding RNA (Lemoine et al. 2013).

We hypothesized that IKKε signaling in corneal epithelial cells in response to TLR stimulation is wired differently from fibroblast or macrophage and serves to stimulate NF-κB activation. As such, we investigated whether K6a and/ or ELKS regulate the phosphorylation and activation statuses of IKKε and RelA. We found that K6a KD hTCEpi cells displayed more stimulatory phosphorylations of IKKε at Ser-172 (Figure 5A, left vs middle panel, lane 3) and RelA at Ser-468 (Figure 5B, left vs middle panel, lane 3) under both basal and PAO1-stimulated conditions. In contrast, the stimulatory phosphorylations of IKKε and RelA in ELKS KD cells were both attenuated (Figures 5A and 5B, left vs right panels, lane 3). These results suggest that K6a suppresses ELKS to promote the phosphorylation and activation of IKKε. We next overexpressed ELKS and IKKε in 293T cells and found that the two proteins steadily interacted with each other as shown in forward and reverse-CoIP experiments (Figures 5C and 5D). Furthermore, K6a and IKKe double KD in hTCEpi cells, mimicking K6a and ELKS double KD, suppressed the increased pro-inflammatory cytokine secretions observed in cells with K6a KD alone under both basal and PAO1-stimulated conditions (Figures 5E, magenta vs cyan bars). Together, these results are in support of a model where K6a antagonizes ELKS-associated IKKε to autophosphorylate and phosphoactivate RelA.

**Figure 5.**
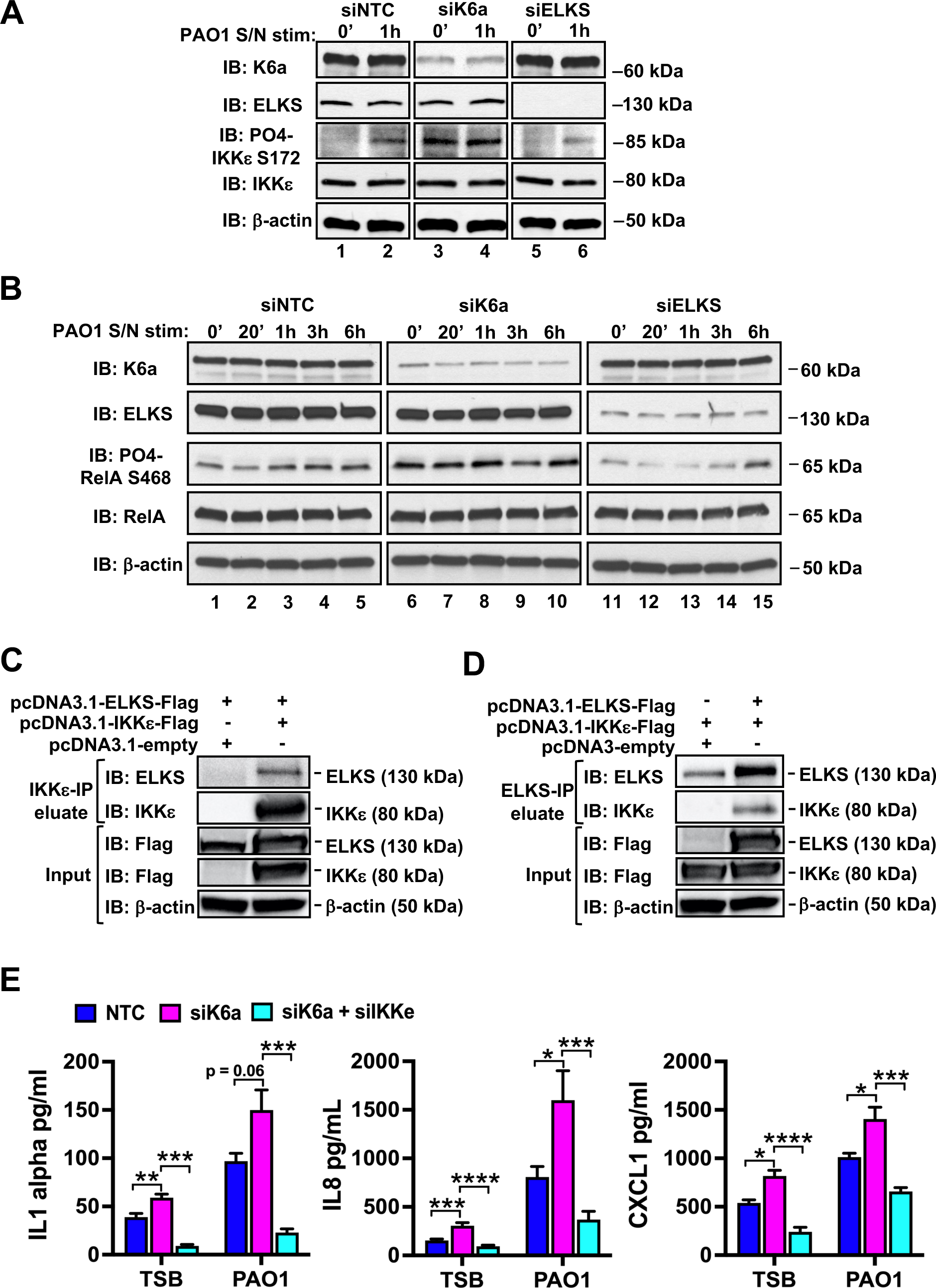
K6a suppresses ELKS-associated atypical IKKε in autophosphorylation and phosphorylation of RelA. **(A)** hTCEpi cells transfected with NTC-, K6a-, or ELKS-siRNAs for 3 days were treated with KGM^TM^-2 media containing 20% (v/v) sterile PAO1 supernatant for 0 min and 1 hour before cell lysis. Equal amounts of cell lysate at each time point were immunoblotted for the detection of K6a, ELKS, PO4-IKKε-Ser-172, total IKKε and β-actin by their respective antibodies. **(B)** Same as (A) except cells were treated for 0 min, 20 min, 1 hour, 3 hours, and 6 hours before cell lysis. Equal amounts of cell lysate were immunoblotted for the detection of K6a, ELKS, PO4-RelA-Ser-468, total RelA and β-actin. **(C and D)** 293T cell lysates containing overexpressed Flag-tagged ELKS and/or Flag-tagged IKKε in combinations indicated were immunoprecipitated with rabbit polyclonal anti-IKKε (C) or anti-ELKS (D) antibodies. Eluted samples were immunoblotted for ELKS and IKKε using their respective antibodies. **(E)** hTCEpi cells transfected with NTC-, K6a-, and IKKε-specific SMARTpool siRNAs as indicated for 3 days were treated with KGM^TM^-2 media containing 20% (v/v) TSB or sterile PAO1 supernatant for 24 hours, followed by conditioned media collection for ELISA. Means ± SEM (n > 3). *P < 0.05, **P < 0.01, ***P < 0.001, ****P <0.0001. Statistical significance was determined by one-way ANOVA followed by Šidák’s post hoc pairwise comparison to NTC- or K6a-siRNA transfected cells within each treatment group.

### K6a knockout C57BL/6 mice display exacerbated inflammation and immunopathology in keratitis models

In the presence of bacterial ligand exposure or infection at the ocular surface, the corneal epithelium quickly produces pro-inflammatory cytokines and chemokines to recruit surrounding myeloid immune cells into the site of challenge and activate their pro-inflammatory and antimicrobial responses. These cells, predominately neutrophil (CD45+ Ly6B.2+ F4/80−), and some macrophage (CD45+ F4/80+), phagocytose and eradiate bacteria by reactive oxygen and nitrogen species attacks (Hazlett 2004; Pearlman et al. 2013; Winterbourn, Kettle, and Hampton 2016). Moreover, they engulf dead cells and secret metalloproteinases to the damaged extracellular matrix, laying the groundwork for tissue regeneration and recovery. However, their lingering activation or clearance at cornea can also induce devastating collateral damage, causing corneal scarring and even blindness (Hazlett 2004; Pearlman et al. 2008; Pearlman et al. 2013).

Because we observed that the corneas of K6a knockout mice produced more cytokines and chemokines than those from wildtype mice early after LPS inoculation (Figure 1), we asked whether their elevated presence has a physiological impact on the subsequent recruitment of myeloid immune cells. We compared between K6a knockout and wild type mice after inoculation of their corneas by purified bacterial ligands (LTA or LPS) for 24 hours in the sterile inflammation model. We isolated their whole corneas and quantified their cytokines and immune cells by ELISA and FACS respectively. In response to LTA, the corneas of K6a knockout mice displayed more robust infiltration of total leukocytes (CD45+), neutrophils (CD45+-Ly6B.2+ F4/80−) and macrophages (CD45+ F4/80+; Figure 6A). Concomitantly, these K6a knockout corneas produced more cytokines (IL-6, GM-SF) and chemokines (CXCL1, CXCL2, CXCL10) than those from wild type mice (Figure 6B). Similar findings were observed in the corneas of K6a knockout mice after LPS inoculation (Figures 7A and 7B).

**Figure 6.**
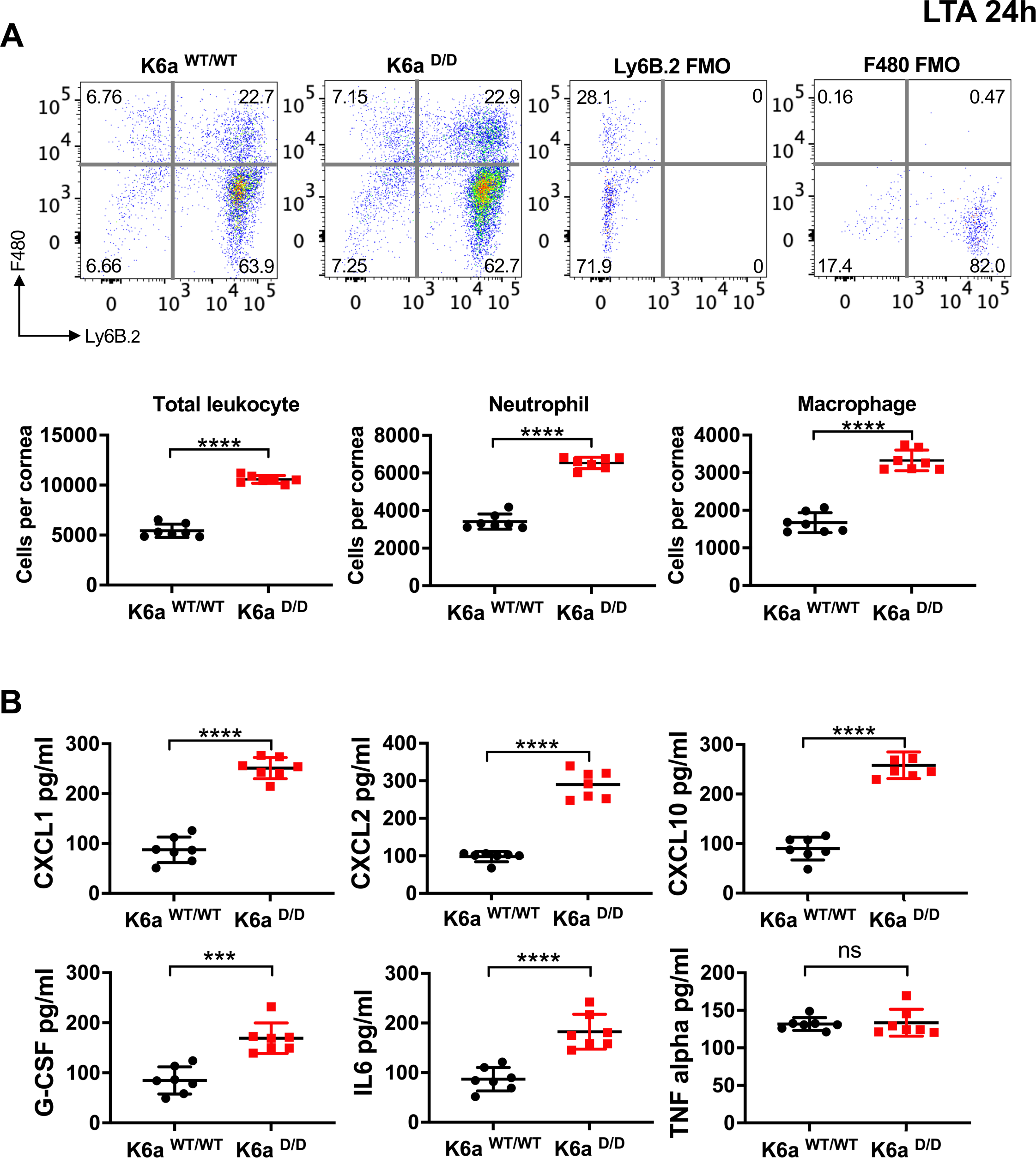
K6a knockout C57BL/6 mice display worsened corneal inflammation in LTA sterile keratitis model. LTA (20μg) was topically inoculated to scarified mouse cornea (one per mouse) of C57BL/6 wildtype (K6a ^WT/WT^) mice and whole body K6a knockout (K6a ^D/D^) mice for 24 hours. Mice were then euthanized and whole corneas were dissected for immune cell and cytokine analysis. Representative fluorescence activated cell sorting (FACS) bivariate dot plots **(A)** showing Ly6B.2- and F4/80-gated cells among total CD45+ cells in the inflamed corneas. Numbers represent percentage of cells in each gate relative to total CD45+ cells. Total leukocytes (CD45+), neutrophils (CD45+Ly6B.2+F4/80−), and macrophages (CD45+F4/80+) per cornea were quantified. ELISA quantification of cytokines per cornea **(B)**. Means ± SD (n = 7 mice (corneas) per group). *P < 0.05, **P < 0.01, ***P < 0.001, and ****P < 0.0001. ns, P > 0.05. Statistical significance was determined by unpaired t test except for CXCL2 which was by Welch’s t test.

**Figure 7.**
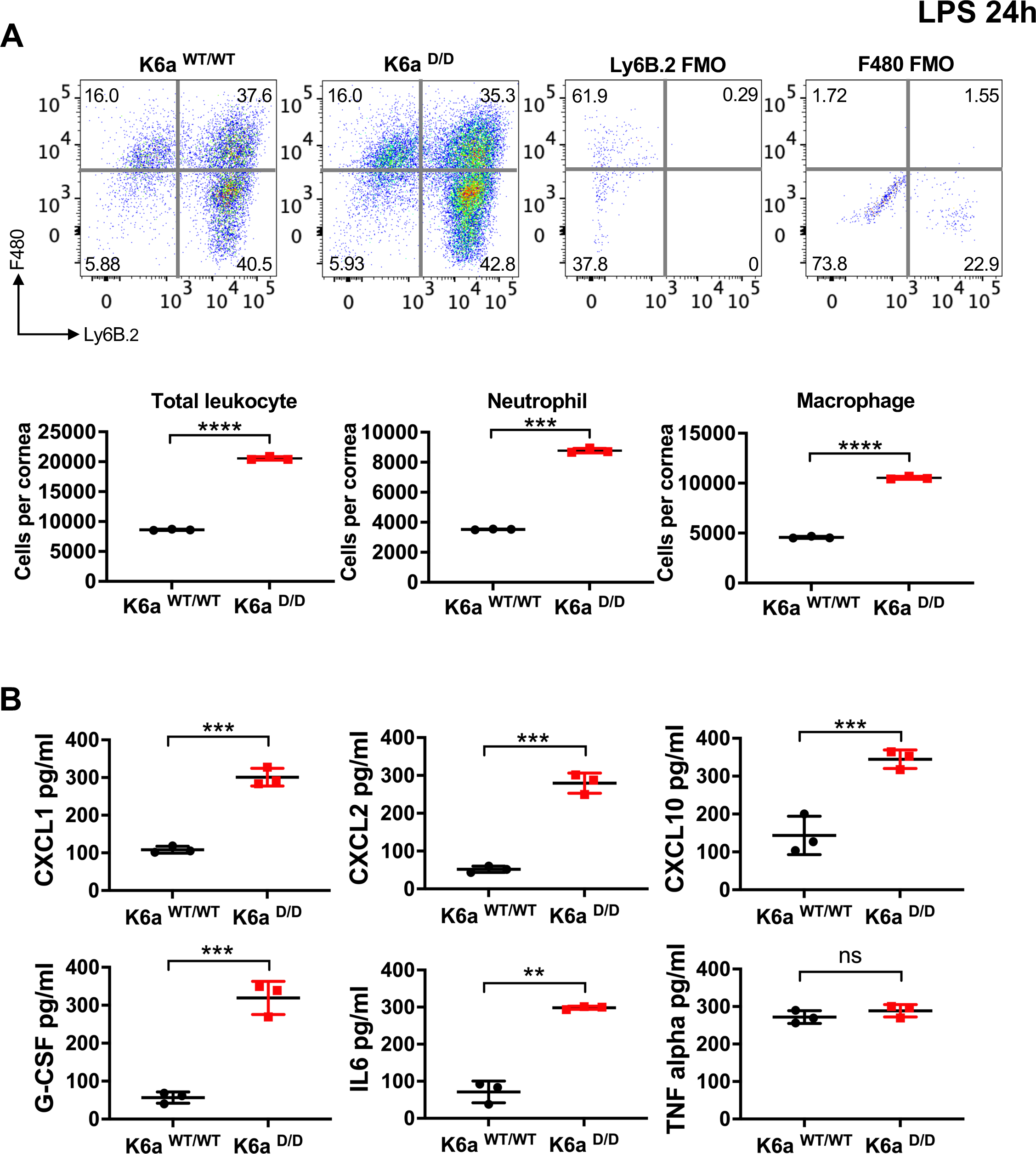
K6a knockout C57BL/6 mice display worsened corneal inflammation in LPS sterile keratitis model. LPS (20μg) was topically inoculated to scarified mouse cornea (one per mouse) of C57BL/6 wildtype (K6a ^WT/WT^) mice and whole body K6a knockout (K6a ^D/D^) mice for 24 hours. Mice were then euthanized and whole corneas were dissected for immune cell and cytokine analysis. Representative fluorescence activated cell sorting (FACS) bivariate dot plots **(A)** showing Ly6B.2- and F4/80-gated cells among total CD45+ cells in the inflamed corneas. Numbers represent percentage of cells in each gate relative to total CD45+ cells. Total leukocytes (CD45+), neutrophils (CD45+Ly6B.2+F4/80−), and macrophages (CD45+F4/80+) per cornea were quantified. ELISA quantification **(B)** of cytokines per cornea. Means ± SD (n = 3 mice (corneas) per group). *P < 0.05, **P < 0.01, ***P < 0.001, and ****P < 0.0001. ns, P > 0.05. Statistical significance was determined by unpaired t test except for IL6 and neutrophil which were by Welch’s t test.

Next, we evaluated the keratitis response in K6a knockout mice and their controls in a bacteria-induced inflammation model where scarified corneas were inoculated with live *S. aureus* bacteria for 24 hours. Here, we employed K6a-conditional knockout mice (K12-Cre ^Tg/null^; K6a ^F/F^) created by the use of transgenic K12-CRE driver mice, further assuring that any changes in corneal innate immunity is caused by specific ablation of K6a expression at the corneal epithelium. We found that the infected corneas of K6a knockout mice developed more opacification than the floxed-K6a control mice (K12-Cre ^null/null^; K6a ^F/F^; Figure 8A) and exhibited a marked increase in disease severity scores (Figure 8B). These corneas also had more robust infiltrated leukocytes (CD45+), neutrophils (CD45+ Ly6B.2+ F4/80−), and macrophages (CD45+ F4/80+; Figure 8D), and produced more cytokines (IL-6, GM-SF) and chemokines (CXCL1, CXCL2, CXCL10; Figure 8E). Similarly, in *P. aeruginosa* keratitis model, the infected corneas of corneal epithelium-specific K6a knockout mice developed more corneal opacities, higher disease severity scores, and produced more cytokines and chemokines (Supplementary figures 3A, 3B and 3D). In both bacteria keratitis models, the corneas of K6a knockout mice also harbored more bacteria; a phenomenon perhaps linked to the loss of KAMPS-dependent antimicrobial activities caused by the knockout (Figure 8C and Supplementary figure 3C).

**Figure 8.**
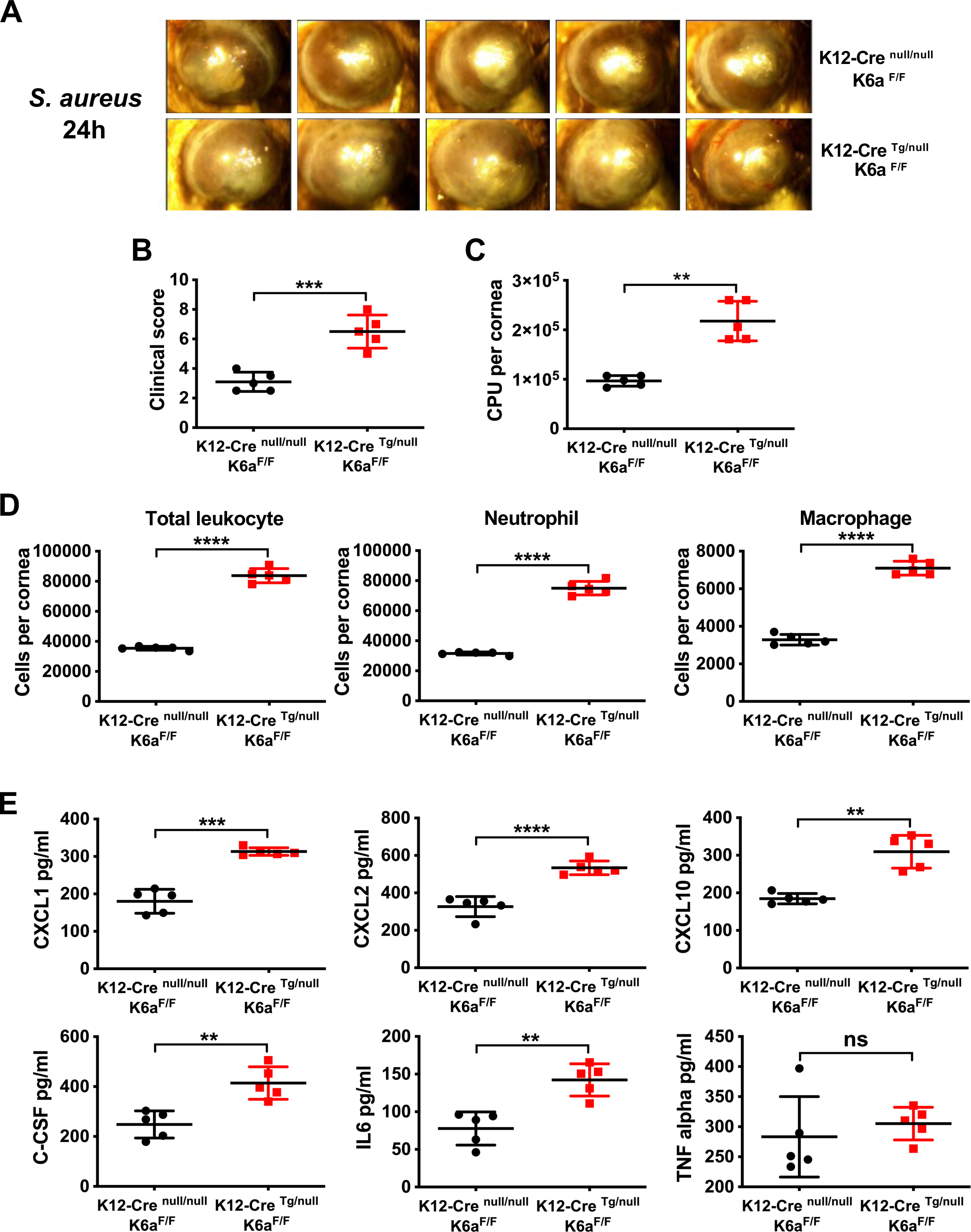
Corneal epithelium-specific K6a knockout mice display worsened corneal pathology and inflammation in S. aureus keratitis model. Scarified mouse corneas (one per mouse) of corneal epithelium-specific K6a knockout mice (K12-Cre ^Tg/null^; K6a ^F/F^) and floxed-K6a control mice (K12-Cre ^null/null^; K6a ^F/F^) were inoculated with *S. aureus* (10^6^ CFU) for 24 hours. Disease presentation **(A)** at 24 hours after inoculation and severity scores **(B)** corresponding to the extents of opacification (area and density) and corneal surface irregularity. Numbers of viable bacteria (**C),** total leukocytes (CD45+), neutrophils (CD45+Ly6B.2+F4/80−), macrophages (CD45+F4/80+) **(D)**, and ELISA quantification of cytokines **(E)** per infected cornea. Means ± SD (n = 5 mice (corneas) per group). *P < 0.05, **P < 0.01, ***P < 0.001, and ****P < 0.0001. ns, P > 0.05. Statistical significance was determined by unpaired t test (clinical score, CXCL2, G-CSF, IL6, TNF alpha, Macrophage) and Welch’s t test (CXCL1, CXCL10, total leukocyte, neutrophil).

Although such higher bacterial burden could fuel some of the immune cell infiltration, given pro-inflammatory cytokine and chemokine productions from LPS-inoculated corneas from K6a knockout mice even pretreated with KAMP10 had remained highly substantial (Figure 1, left black vs right magenta bars), these results overall support the notion that cytosolic K6a’s anti-inflammatory effects at the corneal epithelium during early corneal inflammatory response plays a predominate role in restraining excess myeloid immune cell recruitment, thereby alleviating exacerbated corneal inflammation and immunopathology *in vivo*.

## Discussion

In this work, using K6a knockout mice we have elucidated a novel function of cytosolic K6a in suppressing pro-inflammatory cytokine and chemokine expressions in the corneal epithelium independent of its secreted anti-inflammatory and antimicrobial activities as KAMPs. We previously demonstrated that KAMPS when used alone are highly efficacious in abolishing the ability of myeloid immune cells to activate TLR2/4 signaling (Sun et al. 2023). Specifically, KAMPS interact with TLR2 and TLR4/ MD2 present on their cell surface promoting TLR2/4 endocytosis. The underlying mechanism likely involves the participation of the TLR2/4 co-receptor CD14; another key KAMPS-interacting protein partner (Sun et al. 2023), since ligand-induced CD14 signaling has been proven critical for TLR2/4 endocytosis to occur (Brandt et al. 2013; Roy, Karmakar, and Pearlman 2014; Ciesielska, Matyjek, and Kwiatkowska 2021). Here, using K6a knockout mice we further demonstrated that both KAMPS and cytosolic K6a mediate the suppression of pro-inflammatory cytokine and chemokine productions at the corneal epithelium. Because CD14 is sparsely expressed in the corneal epithelium in vivo (Blais et al. 2005), as opposed to its much higher abundance in myeloid immune cells, the ability of corneal epithelium to employ cytosolic K6a for inflammation suppression could compensate for potential CD14 inefficiency to promote TLR2/4 endocytosis mediated by KAMPS.

Importantly, through proteomic and biochemical analyses, we here further reveal that cytosolic K6a elicits its anti-inflammatory action by serving as an effective antagonist of NF-κB. Previously, the Li group has reported two parallel canonical IKK-dependent pathways downstream of TLR that activate NF-κB (Qin et al. 2006; Yao et al. 2007; Fraczek et al. 2008). The first pathway depends on TAK1 to phosphoactivate IKKα/β, which in turn phosphorylate IκBα to direct its degradation and dissociation from NF-κB. The second pathway relies on MEKK3 to activate IKKα within the IKKα/β heterodimer, which then promotes IκBα phosphorylation and its dissociation from NF-κB without its degradation. As seen here using K6a KD cultured corneal epithelial cells and unlike the first two pathways, we have uncovered a third TLR pathway that does not require any IκBα phosphorylation for NF-κB activation. Instead, it involves a novel interaction between ELKS and IKKε which promotes RelA phosphorylation at serine 468, resulting in enhanced pro-inflammatory cytokine and chemokine expressions. Phospho-RelA (Ser-468) displays longer nuclear occupancy, higher transactivation activities and lower affinity for IκBα as previously reported (Mattioli et al. 2006; Moreno et al. 2010). Because IκBα phosphorylation and degradation require elaborate and concerted actions between kinases and ubiquitin-proteasomal machineries for NF-κB activation, the advantage of this newly described pathway appears that it can quickly deploy RelA for action. Conversely, its downregulation through K6a-mediated suppression of ELKS/IKKε activity does not seem to involve IκBα sequestration or require its resynthesis. In other cell types such as fibroblasts and macrophages, only the canonical IKKs are employed for TLR/NF-κB induction of pro-inflammatory cytokine and chemokine expressions (Hemmi et al. 2004; Perry et al. 2004). Therefore, the use of both canonical and atypical IKKs for NF-κB activation suggests that the cornea epithelium is more flexible at modulating its TLR-inflammatory response *in vivo*, which could greatly minimize corneal damage while preserving its essential functions for barrier protection and light refraction.

Since K6a interacts with ELKS and ELKS KD increases phospho-IKKε (Ser-172), our results suggest that K6a inhibits ELKS by competitive binding against IKKε. While ELKS is an essential scaffold orchestrating IKKα/β-dependent phosphorylation of IκBα and in this study for IKKε phosphoactivation, it remains unclear whether ELKS, NEMO and IKKε assemble into the same complex or they belong to different complexes within corneal epithelial cells. Of note, liquid-liquid phase separation (LLPS) has recently emerged as a key phenomenon driving fundamental biological processes (Mehta and Zhang 2022). In IL-1 receptor-induced canonical NF-κB pathway, NEMO assemble with free and protein-attached K63-linked polyubiquitin chains catalyzed by TRAF6 into LLPS condensates, which are essential for recruiting TAK1 and IKKα/β for their phosphorylation and activation (Xia et al. 2009; Du et al. 2022; DiRusso et al. 2023). Likewise, during IFN-driven interferon-stimulated gene expression, IKKε requires interacting with free K48-linked polyubiquitin chains; an association suggestive of condensate formation, for full activation and STAT1/2-mediated antiviral response (Rajsbaum et al. 2014). Remarkably, ELKS also displays LLPS properties during its regulation of non-NF-κB related cellular processes including vesicle trafficking and pre-synapse organization (McDonald, Fetter, and Shen 2020; Jin et al. 2023). It would be of interest to test whether upon TLR stimulation, the activation of canonical NF-κB and the novel pathway we describe herein involves “phase-association” of ELKS with NEMO or with IKKε into condensates, which in turn serve as distinct signaling activation hubs for their respective pathways. Conversely, K6a may interact with ELKS to exert its anti-inflammatory effect by promoting the dissolution of the interaction between ELKS and IKKε and their associated condensates. Future work will focus on fine-mapping the cytosolic K6a region that attenuates TLR-induced ELKS/IKKε signaling within the corneal epithelium, and test whether this region if over-expressed *in vivo* can alleviate corneal inflammation and immunopathology while promoting speedier recoveries in experimental keratitis models.

Individuals carrying dominant missense mutations at K6 isoforms (K6a, b, c) are predisposed to develop a rare disorder at the skin appendages called pachyonychia congenita (PC). Though not life-threatening, PC patients frequently develop nails dystrophy that are prone to infection and suffer from painful palmoplantar keratoderma (lesions in their palms and soles), greatly impacting on their quality of life. These patients also display other features including oral and hair follicle lesions, excessive sweating and subcutaneous cysts (Leachman et al. 2005; Zieman and Coulombe 2020). The penetrance of PC is low and may require the modulation by additional environmental factors and co-inherited genes (Cooper et al. 2013). In agreement with an earlier report (Wojcik, Bundman, and Roop 2000), we did not detect any PC phenotypes in our K6a knockout mice, lending support to the notion that PC-predisposing K6a mutations are gain-of-function mutations. In contrast, the corneas of these knockout mice markedly increased their cytokine and chemokine productions after being challenged by bacterial ligand as short as 3 hours prior to overt onset of innate immune cell influx. This argues that the anti-inflammatory effect of K6a in the corneal epithelium during the early corneal inflammatory response is a peculiar function that cannot be substituted by other K6 isoforms such as K6b (Takahashi et al. 1998). Meanwhile, an emerging view of TLR function in mucosal epithelium is that it is not merely for antimicrobial defense but is essential for the maintenance of tissue homeostasis, enhancement of barrier integrity, and for the establishment of immunotolerance towards commensal bacteria (Burgueno and Abreu 2020; Fortingo et al. 2022). Intuitively, cytosolic K6a as a TLR antagonist is likely a common strategy employed by K6a-expressing mucosal epithelia such as in the female reproductive tract, oral cavity, tonsils and proximal respiratory tract for both proper resolution and fine-tuning of inflammatory response. In fact, dysregulated epithelial TLR expression or activity has been shown to negatively impact upon the prognosis of HPV positive-related carcinomas and Burkett’s esophageal adenocarcinoma in patients, correlate with their cancer disease progression, and is associated with oral dysbiosis-induced periodontal diseases in experimental models (DeCarlo et al. 2012; Huhta et al. 2016; Jouhi et al. 2017; Delitto et al. 2018). Therefore, the further studying of how cytosolic K6a antagonizes epithelial TLR function could provide new insights on therapeutic approaches that can treat corneal inflammatory diseases as well as other mucosal epithelial diseases for health restoration.

**Supplementary Figure 1. Construction of K6a knockout C57BL/6 mice**

**(A)** C57BL/6 wildtype (K6a ^WT/WT^) mice and whole body K6a knockout (K6a ^D/D^) mice were euthanized. Total RNA was purified and reverse-transcribed from cornea epithelium isolated from dissected cornea. Mouse K6a and GAPDH transcripts were measured by digital droplet PCR. **(B)** Expected genotyping results using genomic DNA extracted from ear tissues from mice that were wildtype (K12-Cre ^null/null^, K6a ^WT/WT^; column 3), homozygous for floxed-K6a (K12-Cre ^null/null^, K6a ^F/F^; column 4), homozygous for transgenic Cre (K12-Cre ^Tg/Tg^, K6a ^WT/WT^; column 5), homozygous for both Cre and deleted K6a alleles (K12-Cre ^Tg/Tg^, K6a ^D/D^; column 6), as well as no template control (column 2). The presence of lox-P sites 1 and 2 flanking the floxed-K6a allele resulted in PCR products larger than the wildtype K6a allele using the same pair of primers.

**Supplementary Figure 2. Effects of K6a knockdown in corneal epithelial cells to cytokine expressions assessed by cytokine antibody array**

hTCEpi cells transfected with NTC- or K6a-specific SMARTpool siRNAs for 3 days were treated with KGM^TM^-2 media containing 20% (v/v) TSB or sterile PAO1 supernatant for 24 hours. Conditioned media were collected and cytokines and chemokines were quantified using the Human XL Cytokine Array kit.

**Supplementary Figure 3. Corneal epithelium-specific K6a knockout mice display worsened bacterial burden in P. aeruginosa keratitis**

Scarified mouse corneas (one per mouse) of corneal epithelium-specific K6a knockout mice (K12-Cre ^Tg/null^; K6a ^F/F^) and floxed-K6 control mice (K12-Cre ^null/null^; K6a ^F/F^) were inoculated with *P. aeruginosa* (10^5^ CFU) for 24 hours. Disease presentation **(A)** at 24 hours after inoculation and severity scores **(B)** corresponding to the extents of opacification (area and density) and corneal surface irregularity. Numbers of viable bacteria (**C)** and ELISA quantification of cytokines **(D)** per infected cornea. Means ± SD (n = 4 mice (corneas) per group). *P < 0.05, **P < 0.01, ***P < 0.001, and ****P < 0.0001. ns, P > 0.05. Statistical significance was determined by unpaired t test.

