## Supplementary figures and images for "Keratin 6a attenuates Toll-like receptor-triggered proinflammatory response in corneal epithelial cells by suppressing ELKS/IKKε-dependent activation of NF-κB"

### Supplemental Figure File

#### Supplementary 1A

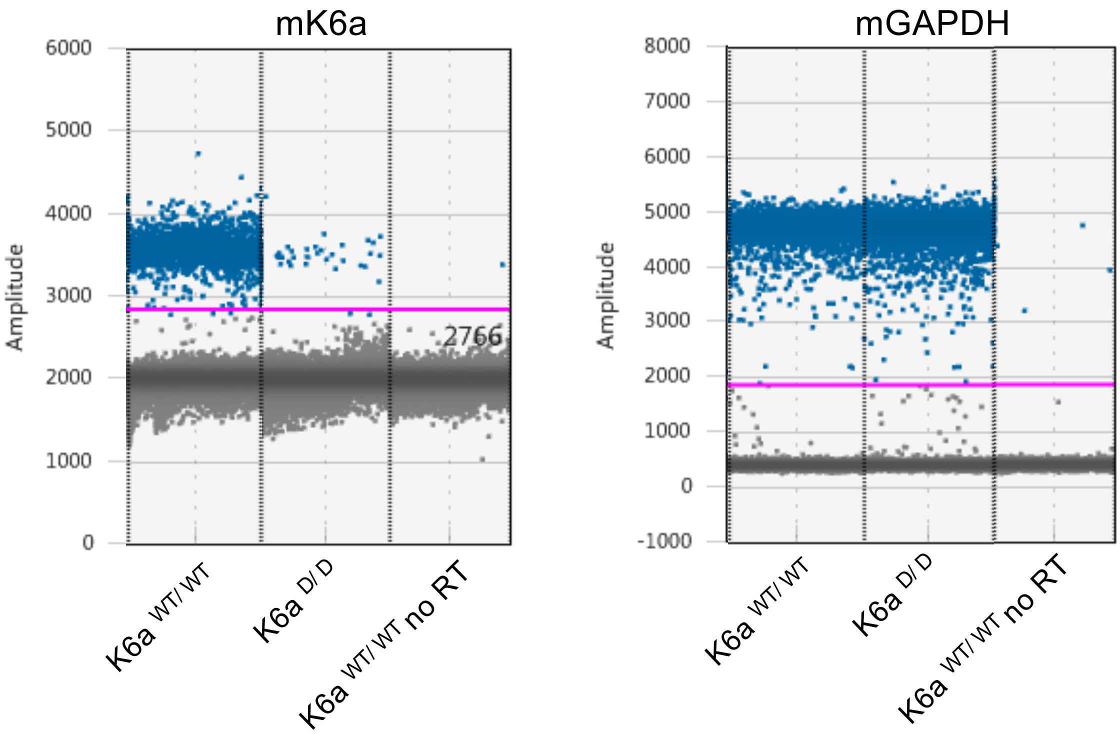


#### Supplementary 1B

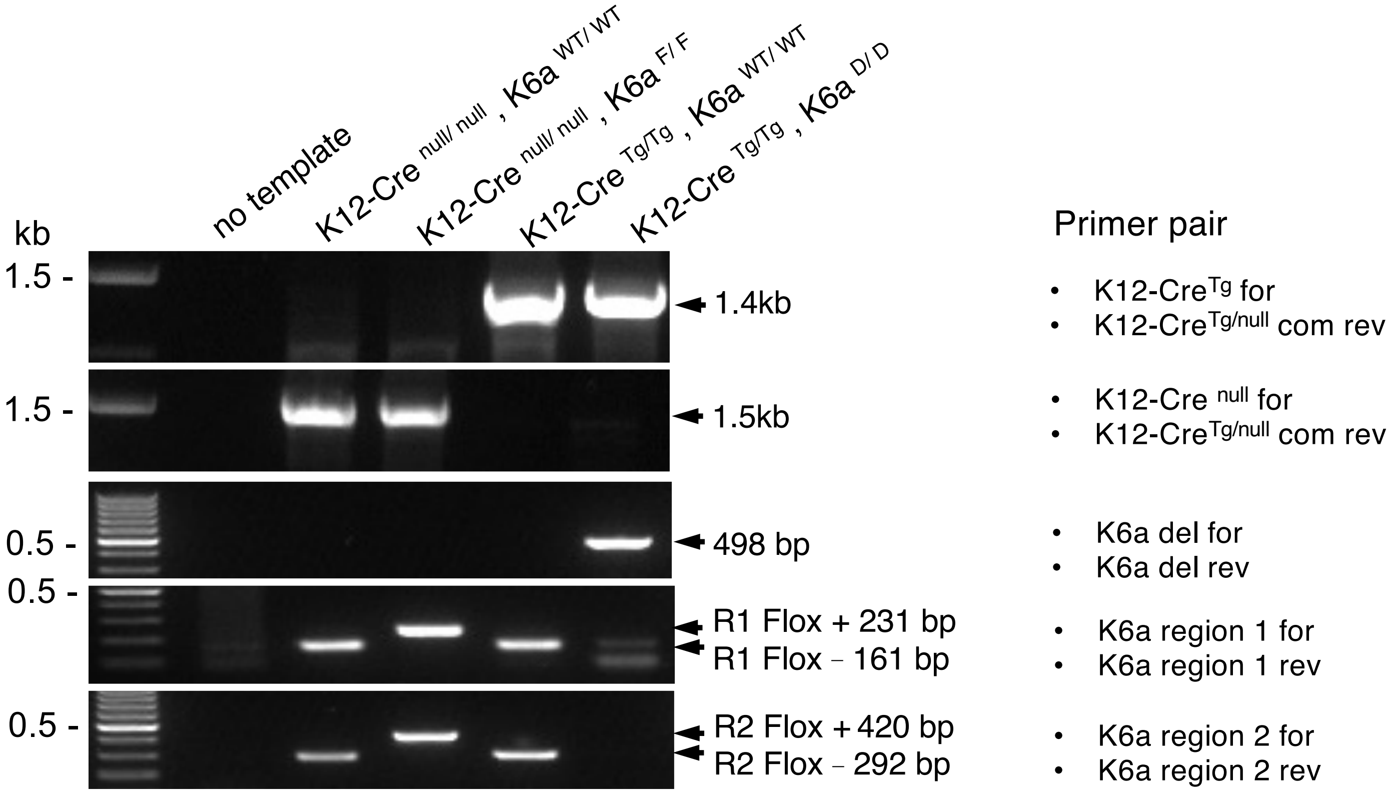

Supplementary 2

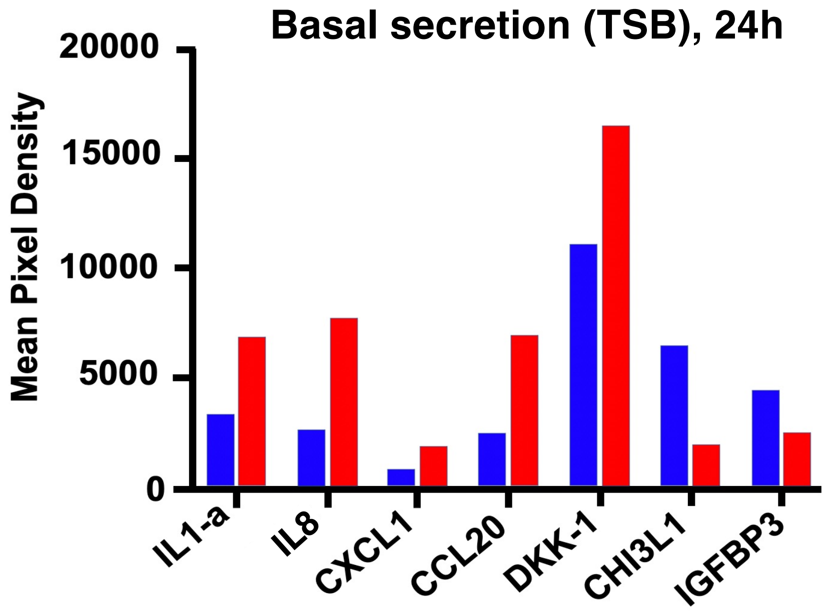


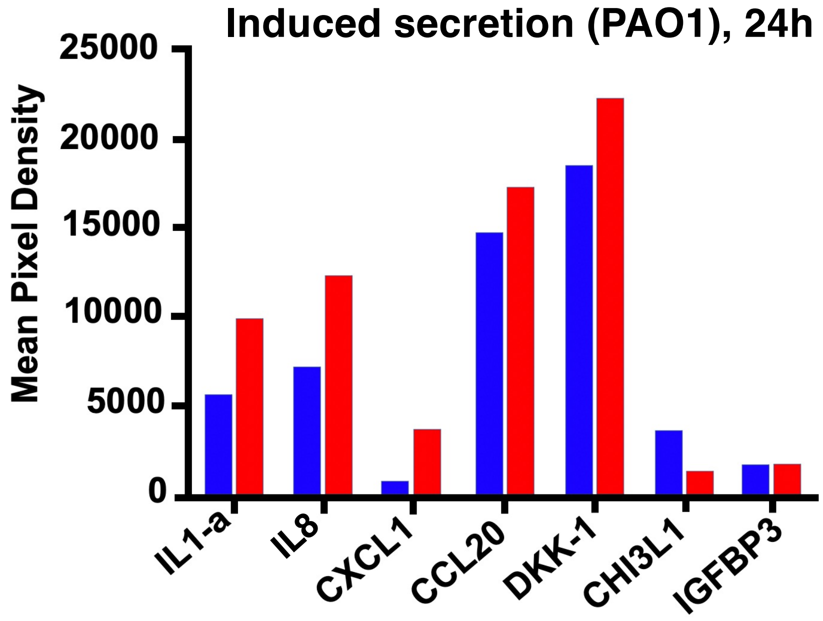

#### Supplementary 3

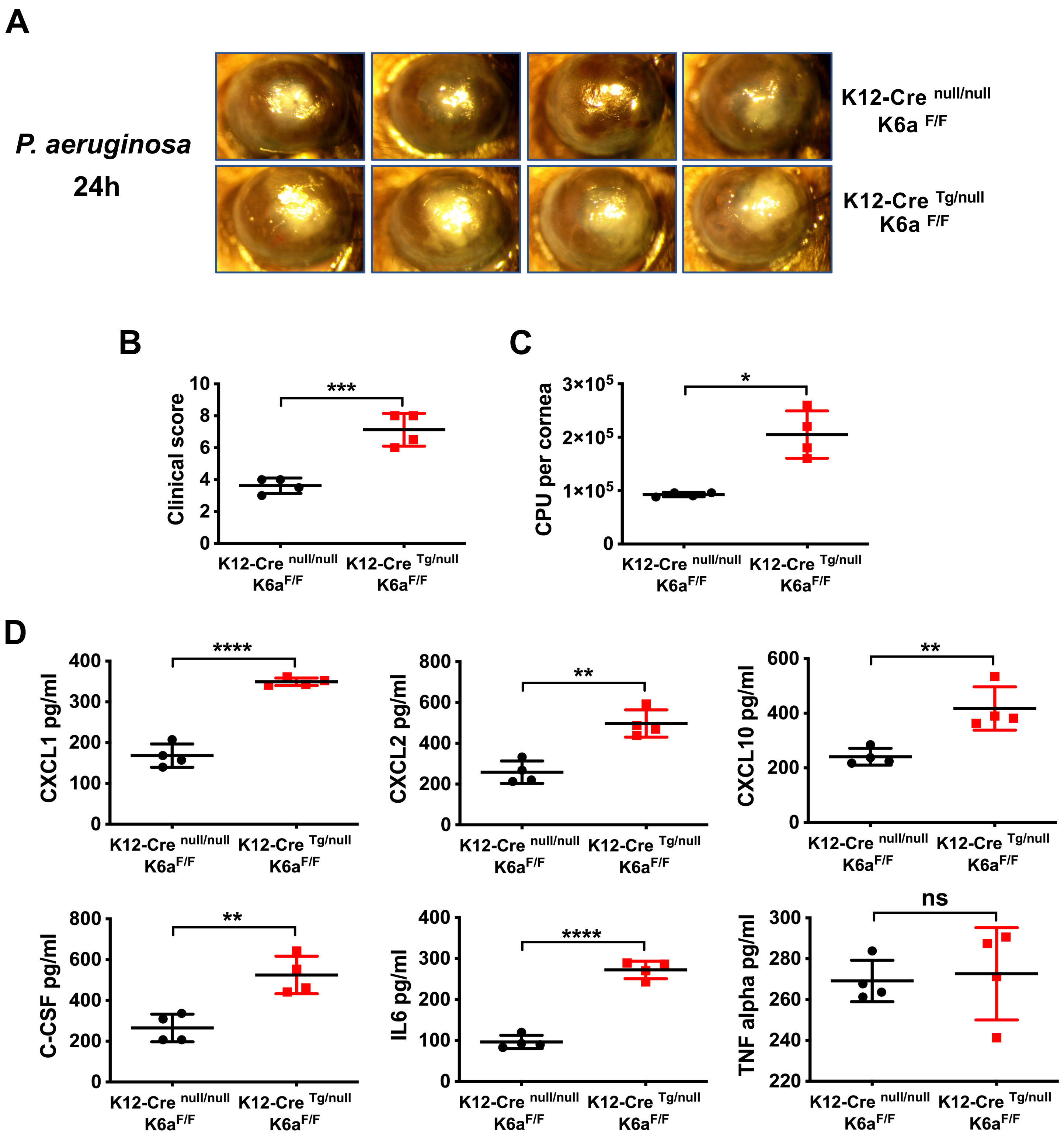
